# A method for partitioning trends in genetic mean and variance to understand breeding practices

**DOI:** 10.1101/2022.01.10.475603

**Authors:** T. P. Oliveira, J. Obšteter, I. Pocrnic, N. Heslot, G. Gorjanc

## Abstract

**Background:** In breeding programmes, the observed genetic change is a sum of the contributions of different groups of individuals. Quantifying these sources of genetic change is essential for identifying the key breeding actions and optimizing breeding programmes. However, it is difficult to disentangle the contribution of individual groups due to the inherent complexity of breeding programmes. Here we extend the previously developed method for partitioning genetic mean by paths of selection to work both with the mean and variance of breeding values.

**Methods:** We first extended the partitioning method to quantify the contribution of different groups to genetic variance assuming breeding values are known. Second, we combined the partitioning method with the Markov Chain Monte Carlo approach to draw samples from the posterior distribution of breeding values and use these samples for computing the point and interval estimates of partitions for the genetic mean and variance. We implemented the method in the `R` package `AlphaPart`. We demonstrated the method with a simulated cattle breeding programme.

**Results:** We showed how to quantify the contribution of different groups of individuals to genetic mean and variance. We showed that the contributions of different selection paths to genetic variance are not necessarily independent. Finally, we observed some limitations of the partitioning method under a misspecified model, suggesting the need for a genomic partitioning method.

**Conclusion:** We presented a partitioning method to quantify sources of change in genetic mean and variance in breeding programmes. The method can help breeders and researchers understand the dynamics in genetic mean and variance in a breeding programme. The developed method for partitioning genetic mean and variance is a powerful method for understanding how different paths of selection interact within a breeding programme and how they can be optimised.

## 1 Background

Analysing genetic trends is essential for identifying key breeding actions and optimizing breeding programmes. The observed genetic change is a sum of contributions from different groups of individuals, often referred to as selection pathways or paths. However, these contributions are difficult to quantify due to the inherent complexity of breeding programmes. The contributions of different groups vary due to different selection intensity, accuracy, genetic variation, generation interval, and dissemination. To quantify the contributions, GarciaCortes et al. [1] developed a method for analysing the change in the genetic mean by partitioning the breeding values into the contributions of several paths. The method uses the standard partitioning of an individual’s breeding value *a*_*k*_ into parent average 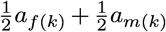and Mendelian sampling term *w*_*k*_. Further, the method assigns Mendelian sampling terms to analyst-defined groups, such as sex, origin, selection path, etc. By aggregating these partitions by other variables, such as year, the method summarises the contributions of different groups of individuals to the overall genetic trend. This approach has been used to quantify the contributions of different countries to the overall genetic trend in the global Brown Swiss population [2], global and local Holstein populations [2], as well as Croatian Simmental cattle [3], and Croatian Landrace and Large-White pigs [4]. More recently, the method was used to estimate the starting point of adopting genomic selection by quantifying differences in genetic trends estimated with pedigree-based and single-step genomic BLUP [5].

In addition to the contribution of paths to changes in genetic mean, breeding programmes should also consider analysing changes in genetic variance to fully understand the drivers of genetic change in a population [6, 7]. Managing the change in genetic mean and variance in breeding programmes is essential to ensure a long-term genetic gain [8, 9]. Therefore, it is crucial that we quantify the contribution of different groups of individuals in a breeding programme to the genetic mean and variance. For example, in several economically important species, male selection and dissemination represent a key lever that has the largest impact on the genetic mean and variance in a population.

The aim of this paper is to extend the method of GarciaCortes et al. [1] to i) partition the genetic mean and variance, ii) implement the method in `AlphaPart R` package, and iii) apply the partitioning method to estimated breeding values following the work of [6] and [7]. We used simulation to demonstrate the methodology and give insights on how to use the `AlphaPart R` package [10] to analyse real data. The remainder of the paper is organised as follows. Section 2 presents the theory of the partitioning method, the extension to partition the genetic variance, and an empirical approach when breeding values are estimated from data. Results are presented in Section 3, where it is shown that the developed approach works well and covers the true values. Generalizations and limitations of the approach are discussed in Section 4, and the conclusions in Section 5.

## 2 Methods

Subsection 2.1 describes the theory of partitioning genetic mean and variance. Estimation of breeding values and partitions from phenotype data with a statistical model and a Markov Chain Monte Carlo (MCMC) method are presented in the subsection 2.2. Subsection 2.3 describes the measures of agreement and bias used to analyse the breeding values and partitions. Subsection 2.4 shows `AlphaPart` implementation working with MCMC samples of breeding values. Next, subsection 2.5 describes simulation of a cattle breeding programme used to demonstrate an `AlphaPart` analysis. Finally, a summary of the used software is described in 2.6.

### 2.1 Partitioning theory

Let ***a*** be a vector of breeding values following a normal distribution with mean **0** and pedigree-based covariance 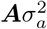 We can write ***a*** as a linear combination of the individual’s ancestor breeding values and individual’s deviation from the ancestors ***a*** = ***Tw***, where ***T*** is a lower-triangular matrix of expected gene flow between ancestors and individuals following a pedigree, and 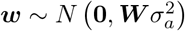 are Mendelian sampling terms representing the deviations, with ***W*** being a diagonal matrix of variance coefficients and 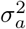 the base population (additive) genetic variance [11, 12, 13, 14].

Assuming a factor with *p* levels, representing our paths (groups) of interest, and for any set 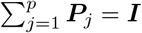 Garcia-Cortes et al. [1] partitioned the gene flow matrix into contributions of each path by defining ***T***_*j*_ = ***T P***_*j*_, *j* = 1, 2, …, *p*, and further partitioned the contribution of each path to breeding values *a priori* using the equality:

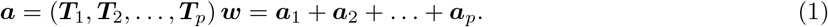

Garcia-Cortes et al. [1] further showed that these contributions can be estimated from data collected in breeding programmes (*a posteriori*). They first calculated the conditional expectation of breeding values given phenotype data (***y***), 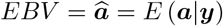. Then they plugged the EBV 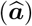 and estimated Mendelian sampling terms 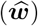 into the equation (1). This approach enabled them to estimate the conditional expectation of partitions, that is, 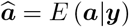:

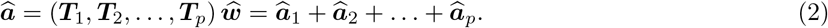

By summarising the breeding values partitions over time, Garcia-Cortes et al. [1] quantified the contribution of each path (for example, males vs. females, different countries, etc.) to the change in individual breeding values and their average over time. Technically this is achieved by sub-setting the 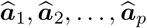 and averaging each subset. This method has been implemented in the `AlphaPart R` package [10, 15]. The `AlphaPart R` package efficiently calculates the partitions by leveraging the sparse ***T*** ^−1^ [11, 12, 13, 14], and enables a straightforward summary by one variable, such as year, or combination of variables (interaction), such as year and sex. The levels of this variable are used to compute the conditional expectation and we refer to it as 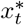, with *t* = 1, 2, …, *m*, and *m* representing the number of distinct categories. Importantly, `AlphaPart` enables use of any function to summarise the partitions of breeding values, that is, *f* (***a***_*j*_).

To enable the use of variance as one of the summary functions in `AlphaPart`, we are here extending the partitioning method to analyse the contribution of paths to genetic variance. Variance of breeding values is, *a priori*, 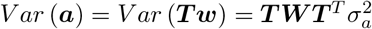 Using the equation (1), we can further partition the genetic variance by paths as:

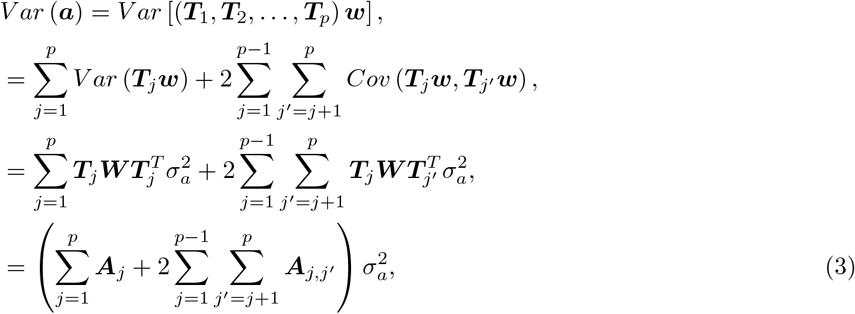

where 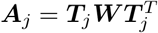 and 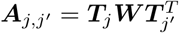 While this partitioning by paths involves dense matrices such as ***A***_*j*_ and ***A***_*j,j*′_, we can efficiently calculate the partitions ***a***_1_, ***a***_2_, …, ***a***_*p*_ by working with the sparse ***T*** ^−1^ [11, 12, 13, 14, 16]. We can again assume 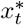 as a variable with *m* distinct categories, *t* = 1, 2, …, *m*, used to summarise the paths. Thus, we can define the genetic variance for the partition *j* given category *t* that has *n*_*k*_ ≤ *nI* individuals, *k*^∗^ = 1, 2, …, *n*_*k*_, as:

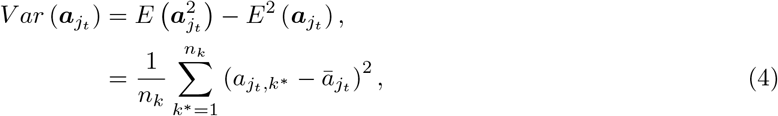

where 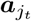is a column for partition *j*, but only considering individuals in category *t, n*_*k*_ is the number of individuals in category *t*, and 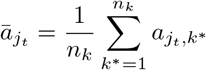. Similarly, genetic covariance between the partitions *j* and *j*^′^, *j*≠ *j*^′^, given category *t* is then:

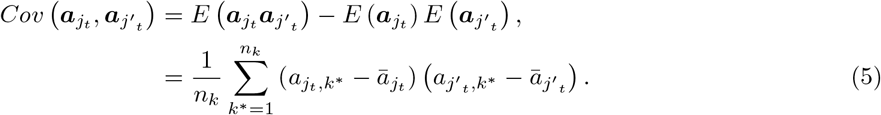

By sub-setting the partitions ***a***_1_, ***a***_2_, …, ***a***_*p*_ by additional variable 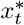 such as year, we can calculate (4) and (5) for each level of the additional variable.

It is worth noting that there is a difference between (3) and (4) or (5). The 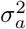 in (3) represents the genetic variance in a base population, while the expression 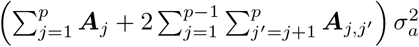 describes how that variance *a priori* changes through a given pedigree and how that variance can be partitioned by paths. Equations (4) and (5) represent the variance and covariance of breeding value partitions ***a***_1_, ***a***_2_, …, ***a***_*p*_, that contribute to the total genetic variance, but conditioned on the category *t* of 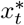, that is, calculated just for individuals in the category *t*. Therefore, we can partition the genetic variance of a population into path contributions, which can be summarised in the same ways as genetic mean [1, 17]. Such analyses can quantify the contribution of different paths to changes in genetic mean and variance over time.

The presented partitioning of genetic variance holds for *a priori* breeding values or true breeding values. However, when EBV are available, we can not substitute ***a***_1_, ***a***_2_, …, ***a***_*p*_ with their expectations 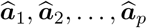 in (4) and (5) as Garcia-Cortes et al. [1] could do it for the partitioning of genetic mean. To see this, imagine a situation where EBV are based on very limited phenotype information. Such EBV will be shrunken strongly towards zero and will have low accuracy [14]. As such, these EBV will not be a good representation of true breeding values, and their variance, *V ar*(*EBV*) = *V ar* (*E* (***a***|***y***)) will be much smaller than variance of true breeding values *V ar* (***a***). To address this issue, we use the approach from Sorensen et al. [6] and Lara et al. [7] that involves three steps. First, sample breeding values from their posterior distribution [16]. Second, for every sample of breeding values calculate desired quantities, in our case breeding value partitions and their means and variances. Multiple samples of these quantities describe their posterior distribution. Third, summarise the samples to describe the posterior distributions of interest, in our case partitions of genetic mean:

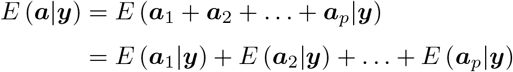

and partitions of genetic variance:

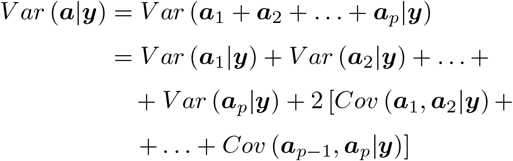

or their variants when we only considering individuals in a specific category.

### 2.2 Statistical model and computational approaches

In the previous subsection, we assumed we knew the true breeding values. However, in reality, we infer breeding values from phenotype data. To this end, we fitted the standard pedigree-based model to observed data:

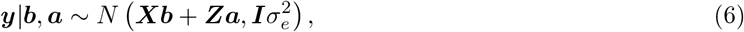

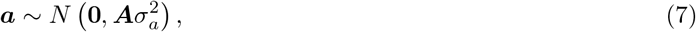

where ***y*** is a vector of observed phenotypes, ***b*** is a vector of fixed effects with the design matrix **X, a** is a vector of breeding values with the design matrix **Z**, 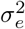 is a residual variance, ***A*** is pedigree-based relationship matrix and 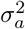 is genetic variance in the base population.

The directed acyclic graph (DAG) representing the model (7) considering only intercept as the fixed effect is illustrated in Figure 1, where pedigree and phenotypic records are displayed in separate plates as a generalization of the case where animals might have a phenotypic record, and the dotted lines indicate a possibly missing parent in the pedigree. In the pedigree plate we have *nI* individuals represented by founders and non-founders, where founder’s breeding values are `a priori` distributed as 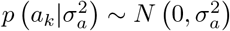. Breeding value of non-founders given their parents are then represented by 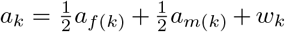, where *a*_*f*(*k*)_ and *a*_*m*(*k*)_ are parent’s breeding values and *w*_*k*_ is the Mendelian sampling term 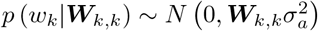.

**Figure 1.**
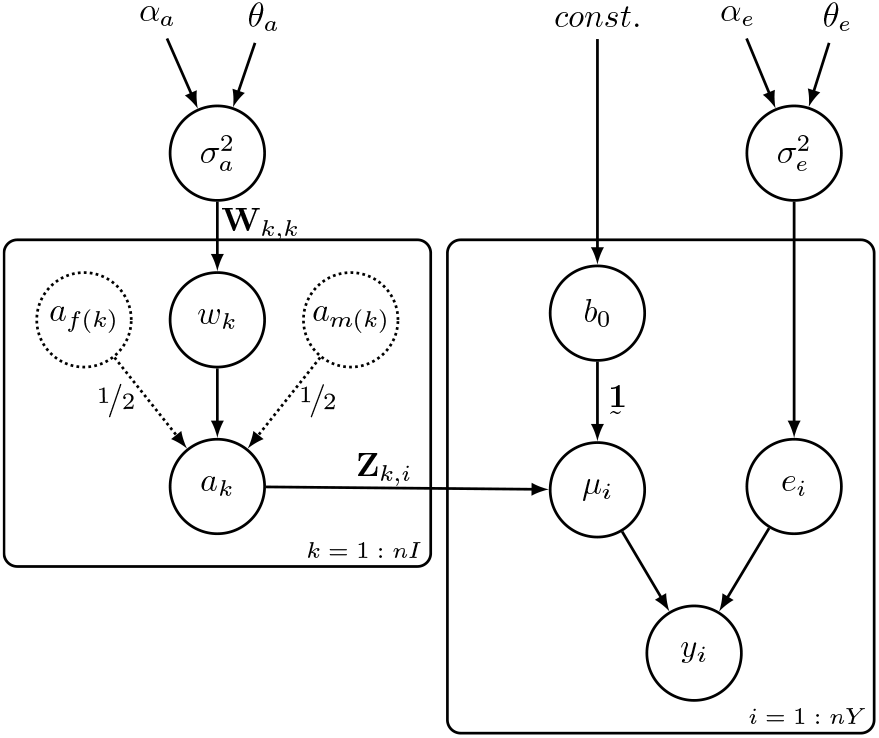
Directed acyclic graph of the pedgiree-based model with *nI* individuals and *nY* phenotypic records (*y*_*i*_) with explicit representation of Mendelian sampling terms (*w*_*k*_) and error term (*e*_*i*_), where 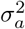 is the additive genetic variance, *a*_*f*(*k*)_ and *a*_*m*(*k*)_ are parent’s breeding value, **1** represents a vector of ones, *µ*_*i*_ the linear predictor, and 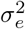 the residual variance

Following equation (3) matrix ***A*** can be decomposed as ***A*** = ***T W T*** ^*T*^ using the generalised Cholesky (LDL) decomposition [11, 12, 13, 14]. The diagonal elements of ***W*** can be computed according to specific scenarios described by [11, 12, 13, 14] as i) 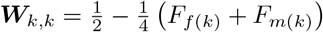 when both parents are known; ii) 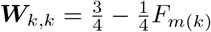 or 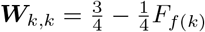 when one parent is known; and iii) ***W***_*k,k*_ = 1 when both parents are unknown, where *F*_*f*(*k*)_ and *F*_*m*(*k*)_ are respectively the coefficients of inbreeding of the father and mother of the individual *k* [11, 12, 13, 14]. While this theory is standard, some quantitative genetics software still ignore parental inbreeding coefficients in setting up the ***A***^−1^, which can have significant impact on the analysis of genetic variance as we will show in results.

We used the full Bayesian approach to infer breeding values using (7). To this end we specified prior distribution for all model parameters, as shown in Figure 1. Thus, ***b***, 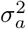, and 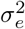 were assumed to have a joint prior density of the form 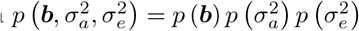 with a flat distribution for *b* and a conjugate inverse-gamma(*α, θ*) distribution for variances (or gamma distribution for precisions = 1/variance), where *α* and *θ* are set to a value such as 0.1^3^:

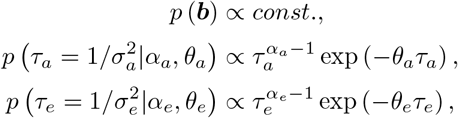

with *τ*_*a*_ *>* 0, *α*_*a*_ ≥ 0, *θ*_*a*_ ≥ 0, *τ*_*e*_ *>* 0, *α*_*e*_ ≥ 0 and *θ*_*e*_ ≥ 0. The posterior distribution is then obtained by applying

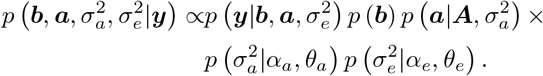

the Bayes’ theorem conditional on the data:

We sampled from the posterior distribution with a Markov Chain Monte Carlo (MCMC) algorithm (the Gibbs algorithm) as implemented in [18]. We constructed one chain with 80,000 samples, from which 20,000 were considered as burn-in, while the remaining 60,000 were stored and thinned by saving every 40-th sample. We assessed the MCMC convergence by inspecting the trace and auto-correlation plots. Consequently, 1,500 samples of breeding values were stored representing the posterior distribution *p* (***a***|***y***). These samples were passed as input to the `AlphaPart R` package.

It is imperative to note that the proposed partitioning method requires samples from the posterior distribution 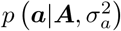 to enable inference of the path contributions to both genetic mean and variance. While we have used the full Bayesian approach with MCMC [16], an alternative is to use the empirical Bayesian approach, that is, estimating variance components with REML and sampling breeding values assuming that variance components are known (see [7]). Obviously, the full Bayesian approach is recommended to account for uncertainty in estimating all model parameters.

### 2.3 Measures of agreement and bias

The partitioning methodology depends on unbiased estimates of breeding values. If the used model (7) does not adequately describe the data, we can end up with biased inference of ***a***. Let *t* be the index for generation with *t* = 1, 2, …, *m*. In simulations we can calculate *E* (***a***_*t*_), *E* (***a***_*t*_|***y***), *V ar* (***a***_*t*_), and *V ar* (***a***_*t*_|***y***), where ***a***_*t*_ and ***a***_*t*_|***y*** are respectively vectors containing the true and inferred breeding values of all individuals from generation *t*. To evaluate unbiasedness of breeding values, we can evaluate the agreement between 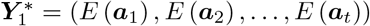 and 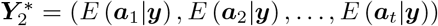. Note that the same holds if we are interested in the agreement between variances, we only need to replace *E*(.) by *V ar*(.). The same holds for breeding value partitions.

Let us assume that the pairs of 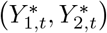 are independent draws from a bi-variate population with means *µ*_1_ and *µ*_2_ and a covariance matrix:

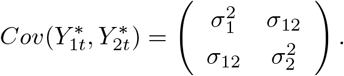

We can evaluate the agreement between 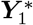 and 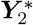 with the concordance correlation coefficient [19], which is an agreement index that lies between −1 and 1, and is given by:

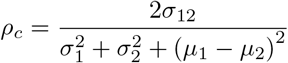

where 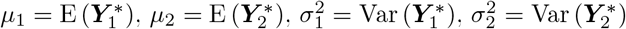 and 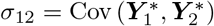. It can be shown that *ρ*_*c*_ = *ρ* × *C*_*b*_, where *ρ* is the Pearson correlation coefficient (a measure of precision), and *C*_*b*_ is the bias correction factor (a measure of accuracy). Here, *ρ* measures how far each observation deviates from the best-fit line, and *C*_*b*_ ∈ [0, 1] measures how far the best-fit line deviates from the identity line *y* = *x* and is defined as *C* = 2 (*v* + *v*^−1^ + *u*^2^)^−1^, where 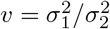 is a scale shift and 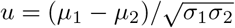 is a location shift relative to the scale. When *C*_*b*_ = 1, there is no deviation from the identity line. In this sense, 1 − *C*_*b*_ can be seen as a measure of bias.

We can define bias as a measure of the systematic error of an estimator by evaluating the difference between the expected value of a statistic and the true value of its corresponding parameter. In this study, we also want to investigate the distribution of the difference between true and inferred quantities of interest. Our quantities of interest were partitions of genetic mean and variance over various categories. Let *n*_*r*_ be the number of simulation replicates, *r* = 1, 2, …, *n*_*r*_, and 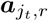 and 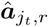respectively, the *j*-th true and inferred partitions of breeding values for the category *t* at replicate *r*. As we are using a Bayesian approach, we can obtain the posterior distribution of our quantities of interest, 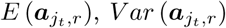, and 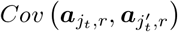. We summarised these posterior distributions with the posterior mean and calculated the difference 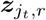between this posterior mean and the true value for means, variances, or covariances. Thus, we can assume the difference 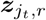 follows a normal distribution for a given partition *j* and category *t*:

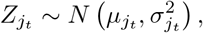

where 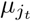and 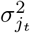 represent the mean and variance of 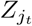. Specifically, we inspected the distribution of the difference 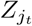for each partition *j* and generations as categories indexed by *t*.

### 2.4 AlphaPart Implementation

We implemented the partitioning method in the `AlphaPart R` package [10, 17] with the summary method:

~~~
R> summary(object = part, by = “time”, FUN = var)
~~~

where `object` is an object of class `AlphaPart` with path-partitioned breeding values, `by` represents the name of a column by which summary function `FUN` should be applied; and `FUN` is the summarising function. We also included an extra argument called `cov` that controls how the covariances will be displayed in the output. If `cov = FALSE`, the default, all covariances will be returned in a single column represented by 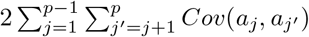, otherwise, if `cov = TRUE`, the `summary` method returns *p* (*p* − 1) */*2 columns, where each column represents covariances in the form of 2*Cov*(*a*_*j*_, *a*_*j*′_).

We further describe how to use the posterior samples of breeding values from subsection 2.2 in `AlphaPart`. Let *T* be the number of traits and *S* be the number of samples of breeding values. Suppose `data` is a data frame containing columns for individual (`id`), father (`Fid`), mother (`Mid`), path (`path`), and generation (`Gen`). Now suppose a more general case where `bv_samples` represents a data frame containing a column for the individual (`id`) identification and *S* columns for the samples of breeding values, as shown in Figure 2. To prepare the input data for `AlphaPart`, we can merge the data frames called `data` and `bv_samples` into a new data frame called `newData` (Figure 2). We can then use the function `AlphaPart()` to calculate breeding value partitions with the difference that we now should pass the samples names to the argument `colBV` (Figure 2). Afterwards, the `summary()` function can be called to summarize the partitions using an explanatory variable, such as generation (`Gen`). Since we work with posterior samples of breeding values we obtain posterior samples for the summaries of the partitions (see the accompanying code). Finally, in the case with more than one trait, we suggest a `for` loop (possibly parallelized) to create one output per trait, as shown in Figure 2. In an extreme case with more traits than samples an alternative approach would be to save one sample of breeding values for multiple traits in one data frame and loop over the samples.

**Figure 2.**
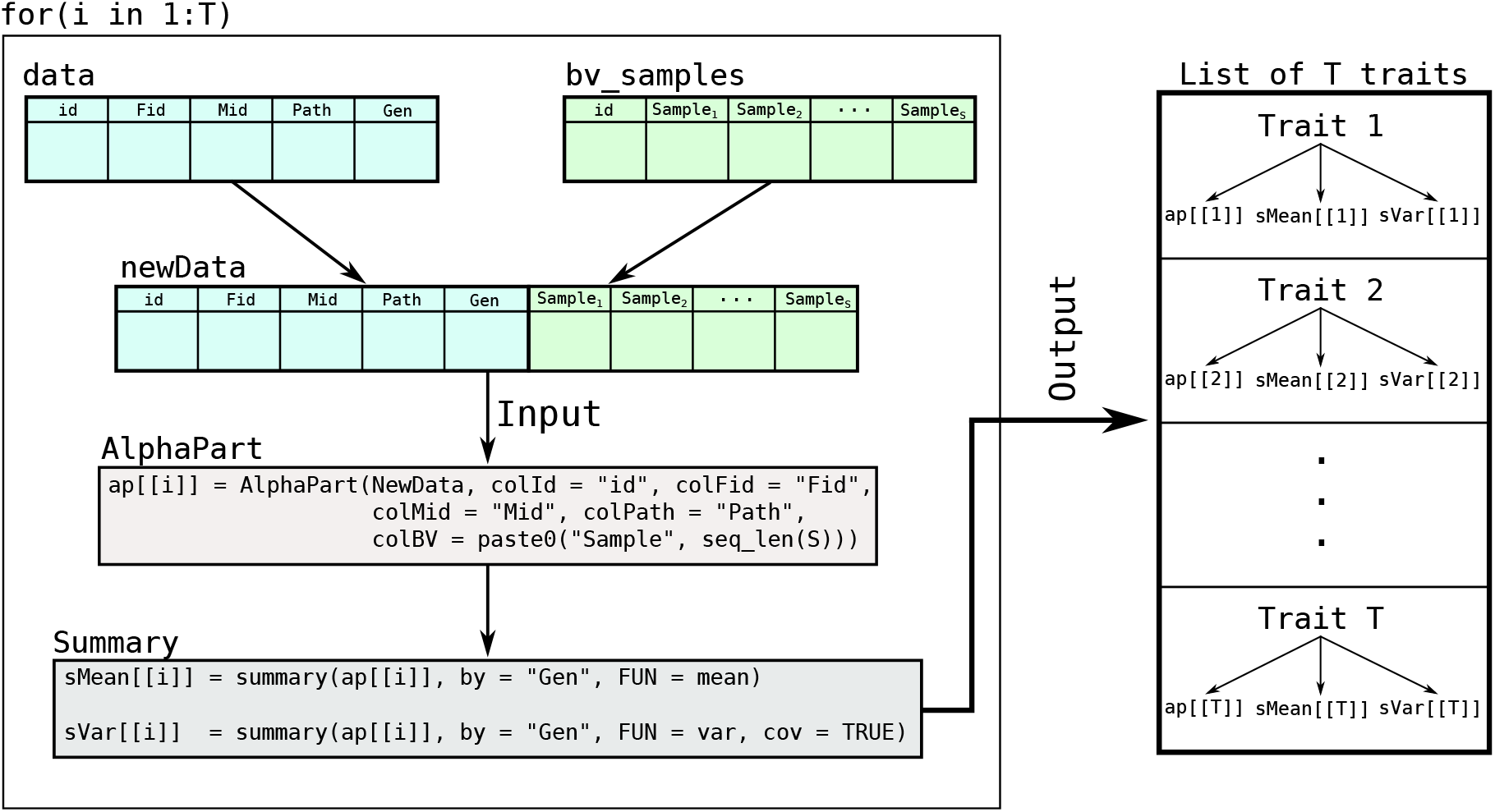
Flowchart representing a possible algorithm to evaluate contributions of paths to genetic mean and variance using samples of breeding values with the `AlphaPart R` package in a multi-trait case

### 2.5 Simulation

To evaluate the method and `AlphaPart` implementation, we used `AlphaSimR` to simulate a simple cattle breeding programme over 40 generations with 1,000 individuals per generation. The first 20 years represented a burn-in phase, where we selected the best 5 males (out of 500) as sires based on their phenotype and mated them with all 500 females from the previous generation and all 500 females from the current generation. These matings produced 1,000 selection candidates for the next generation. After the burn-in phase, we tested two selection scenarios over further 20 generations: we selected 5 best males from 500 male candidates based on i) their phenotypes (‘medium-accuracy’ scenario, *r* = 0.3) or ii) true breeding values (‘high-accuracy’ scenario, *r* = 1), as shown in Figure 3. We replicated the simulation 30 times with the same founding genomes.

**Figure 3.**
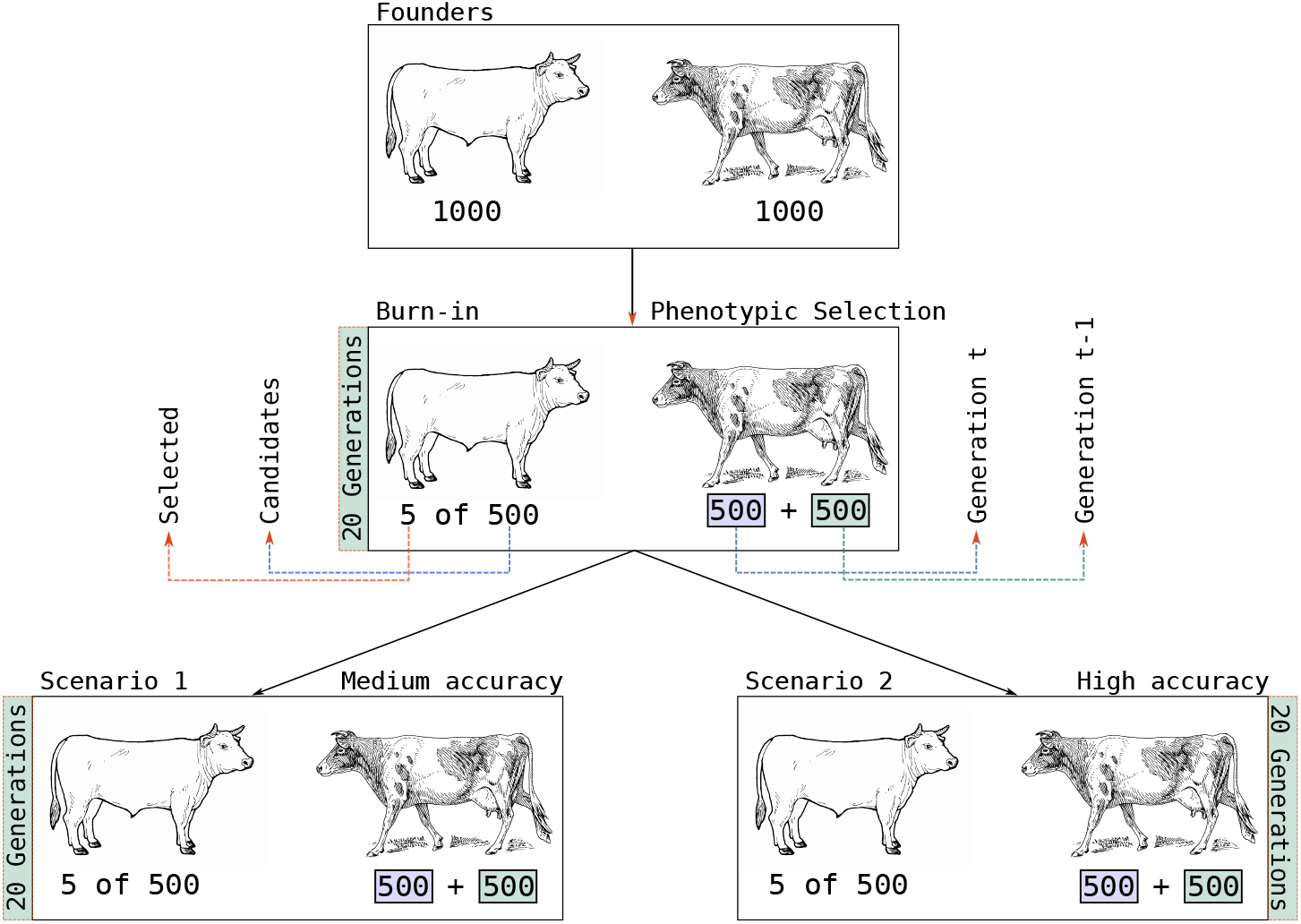
Simulation scheme illustrating an overview of the medium- and high-accuracy scenarios

The simulation was done with the `AlphaSimR R` package version 1.0 [20]. We simulated a cattle genome with 30 chromosomes, 30,000 quantitative trait loci (QTL) with additive effects sampled from a normal distribution, single-trait phenotype with a heritability of 0.3. The above described breeding programme has a low effective population size (for example, *N*_*e*_ ∼ (4 × *nSires* × *nDams*)*/*(*nSires* + *nDams*) = (4 × 5 × 1000)*/*(1005) *<* 20, because we aimed to generate an intense selection situation that would show changes in genetic mean and variance. We split the gene-flow matrix ***T*** by specifying male and female paths (***P***_*m*_ + ***P***_*f*_ = ***I***) and further splitting the male path into selected and non-selected path 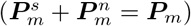, where ***P***_*m*_ is a diagonal matrix with ones in rows for males and zeros otherwise; 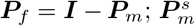 is a diagonal matrix with ones in rows for selected males, and 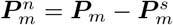 is a diagonal matrix with ones in rows for non-selected males.

### 2.6 Software implementation

We have simulated the cattle breeding programme using `AlphaSimR R` package [20]. We have fitted the model (7) using `BLUPF90` family of programs [18] while all post-processing operations were done in `R` [15]. To compute and summarise the partitions we used the `AlphaPart R` package [10], to prepare data and present results we used the collection of `tidyverse R` packages [21] and `patchwork R` package [22]. The simulation and analysis code is fully available at the GitHub repository https://github.com/HighlanderLab/toliveira_alphapart_variance.

## 3 Results

We used a simulated cattle breeding programme to illustrate the method and software implementation and divided the results into two subsections. Subsection 3.1 describes the genetic mean and variance over generations for both medium- and high-accuracy scenarios and summarises the partitioning results using true breeding values. In the subsection 3.2 we show partitioning results when breeding values are estimated from phenotype data, comparing the estimated and true partitions.

### 3.1 Partitioning of true breeding values

Analysing true breeding values is a crucial way to demonstrate how the partitioning of breeding values and their means and variances works without the uncertainty due to estimating breeding values. Figure 4A shows partitions of genetic mean and variance by sex (males and females), while Figure 4B represents partitions by sex and selection status (selected males, non-selected males, and females) over generations. We scaled the genetic mean and variance of the base population respectively to zero and one to facilitate interpretation. We note that while we partitioned true breeding values, simulation was driven by selection with medium or high accuracy. This means that the accuracy impacted true trends in genetic mean and variance, and we analysed these simulation outputs.

**Figure 4.**
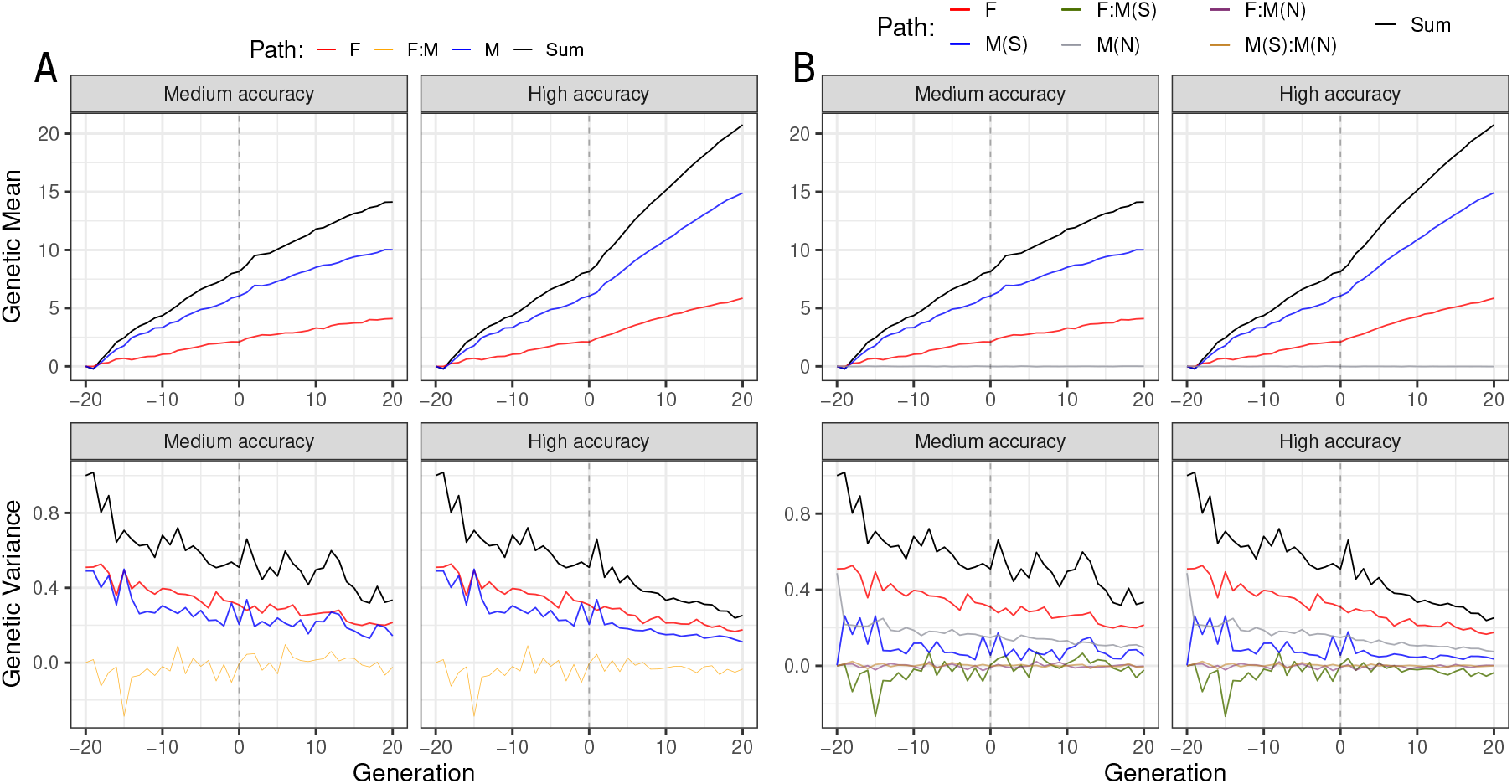
Partitions of genetic mean and variance by (A) sex (males and females) and (B) by sex and selection status (selected males (M(S)), non-selected males (M(N)), and females (F)) using true breeding values for one simulation replicate

Figure 4A shows that males contributed almost twice as much to the genetic mean as females. Interestingly, females were not selected at all (all females contributed progeny for two generations), so they contributed to genetic gain through the dissemination of the past male selections as we will show below. The higher accuracy expectedly drove larger changes in genetic mean and variance than the medium accuracy. If we would use only sex to analyze the drivers of change in genetic variance, we would have concluded that both male and female partitions contributed similarly to genetic variance for both selection scenarios. This observation raises a question: “How can males and females contribute a similar proportion to the genetic variance over time, if we are selecting among males, but not among females?”. The answer to this question is shown in Figure 4B where we partitioned breeding values by sex and selection status. Although non-selected males did not contribute to the change in the genetic mean, their partition still contributed to the genetic variance in their generation. As expected 5 selected males contributed the least to genetic variance, followed by the 495 non-selected males and 500 females (born in a specific generation). Furthermore, since the male partition encompasses selected and non-selected males, this partition spans a wide distribution of partitioned breeding values and this wide distribution has a large variance. This is shown in the Supplementary Figures S1 and S2 with distributions of breeding value partitions for all 40 generations.

Comparing the medium- and high-accuracy scenarios showed that the contribution of paths to genetic variance is a function of selection accuracy, with higher accuracy driving more changes in genetic variance. Notably, with medium accuracy, we saw a smaller difference between partitions of genetic variance for selected and non-selected males. The main reason for this is that the medium accuracy likely did not enable the selection of the top males from the tail of the distribution, which would have much lower variance.

Splitting the male path into selected and non-selected paths also showed that the negative covariance between male and female partitions in Figure 4A was driven by the covariance between female and selected male partitions (F:M(S), Figure 4B). This covariance was consistently negative from generation 8 to 20 in the high-accuracy scenario (Figure 4B), resulting in a mean correlation of −0.33 (±0.15) for that generation interval. As a result, the total genetic variance in a generation *t* can be smaller than the sum of genetic variances for partitions. This non-independence of partitions of genetic variance is more evident in the high-accuracy scenario from generations 8 to 20, where the correlation decreased even more than in the medium-accuracy scenario (Supplementary Figure S3). The non-independence of partitions of genetic variance is yet another reason why individual partitions of genetic variance have to be interpreted with caution.

To further clarify why female partition had a non-zero contribution to the genetic gain, despite the absence of selection among females, the Figure 5 shows the distributions of breeding value partitions and Mendelian sampling terms by the group in generation 39 for medium-accuracy (Figure 5A) and high-accuracy (Figure 5B) scenarios. We can see that the female partition contributed significantly to genetic gain (red distribution), though clearly less than the selected males partition (blue distribution) and clearly more than the non-selected males partition (gray distribution). The inspection of Mendelian sampling terms for females, and for non-selected males, expectedly showed no selection - their Mendelian sampling terms were distributed around zero, while selected males had consistently positive Mendelian sampling terms. However, females were the progeny of previously selected males, and their sons were subject to selection, which creates a non-zero contribution to the female partition - through the dissemination of genes selected in their sires and through their (dam’s) sons.

**Figure 5.**
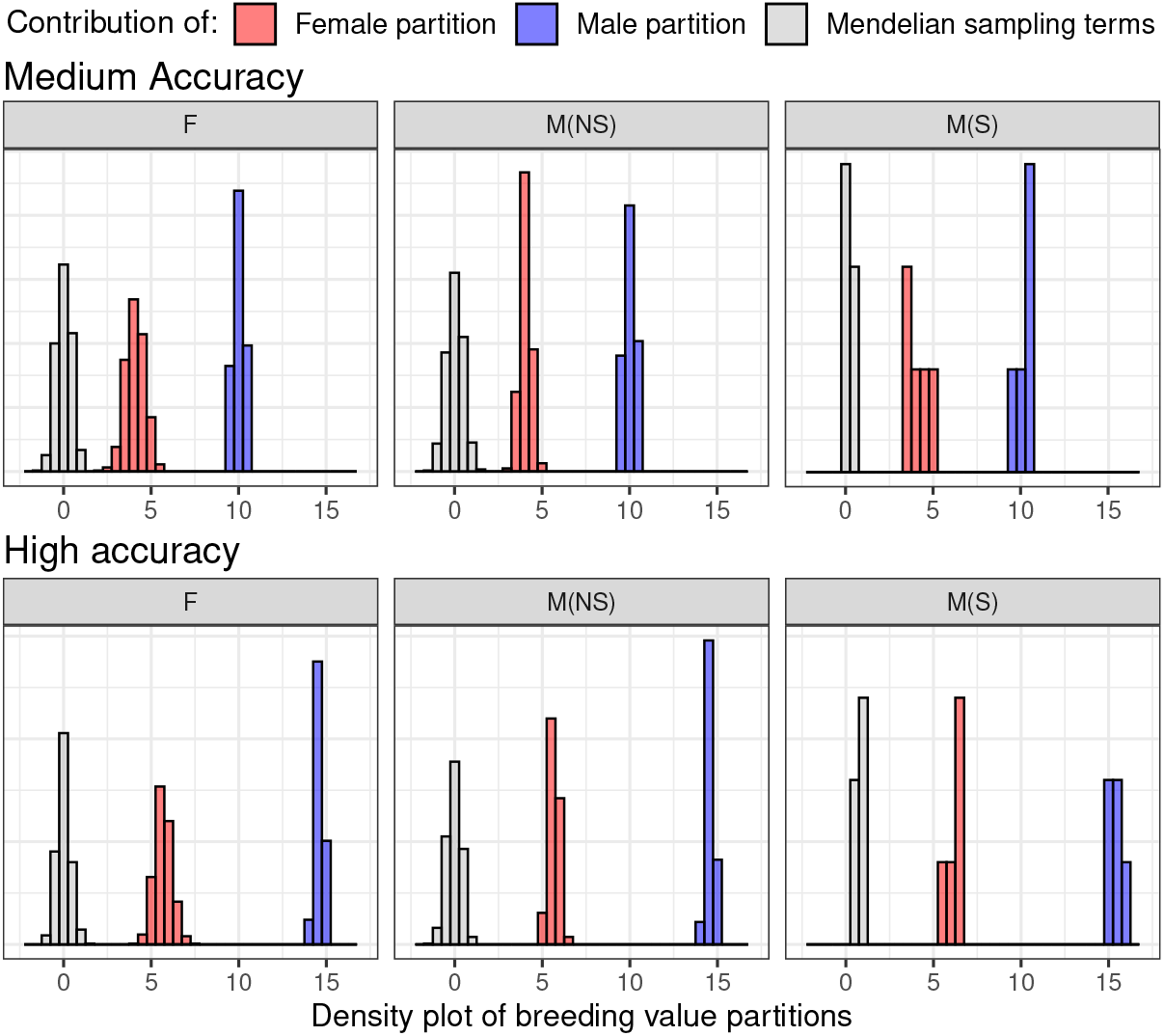
Distribution of breeding value partitions and Mendelian sampling terms by group in generation 39 for medium-accuracy (A) and high accuracy (B) selection scenarios

The presented results showed one replicate of the simulation. In the supplementary Figure S4 we show the partitioning analysis for each of the 30 replicates that all used the same founding genomes. Our aim was to show that the above results are consistently observed across many replicates, but to also show the magnitude of variation between replicates. The Supplementary Figure S4 shows the 30 replicates with the solid line representing the median and the ribbon represents the distribution of true partitions of genetic mean and variance as well as correlation between selected male and female partitions.

### 3.2 Estimating the partitions of genetic mean and variance

#### 3.2.1 Model fit

The data were analysed with the model (7) using the complete pedigree that enabled accurate estimation of residual and base population genetic variance, though we slightly overestimated base population genetic variance in the high-accuracy scenario (Table 1). Evaluating the model further in the terms of estimating the quantities of interest well, we observed good agreement and unbiased estimates under the medium accuracy for genetic means over generations and slightly lower agreement and some bias for the genetic variance over generations (Table 2). Under the high accuracy, the genetic mean over generations was also well estimated, but there was considerable disagreement and bias for the genetic variance over generations (Table 2). The estimated and true genetic means and variances over 40 generations are shown in Supplementary Figures S5 and S6. One reason for a worse performance of the model (7) under the high accuracy, was that this scenario generated significant genetic change both in mean and variance (Figure 4, which was also manifested in a higher level of inbreeding than the medium-accuracy scenario (Supplementary Figure S7). As inbreeding increases over generations, it generates variation between individuals that is challenging to represent using only pedigree-based relationships and better approaches are needed, such as, genomic relationships.

**Table 1.**
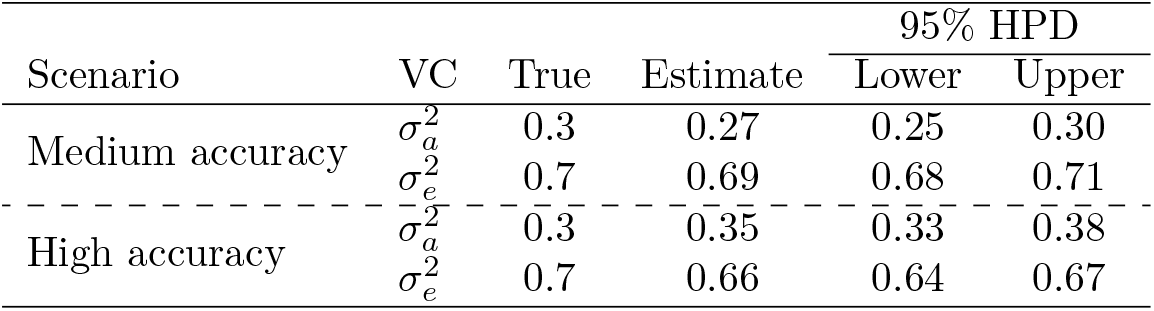
Variance components (VC) true values, point estimates, and their 95% highest posterior density (HPD) interval

| Scenario | VC | True | Estimate | 95% HPD |  |
| --- | --- | --- | --- | --- | --- |
|  |  |  |  | Lower | Upper |
| Medium accuracy | $\sigma_a^2$ | 0.3 | 0.27 | 0.25 | 0.30 |
| | $\sigma_e^2$ | 0.7 | 0.69 | 0.68 | 0.71 |
| High accuracy | $\sigma_a^2$ | 0.3 | 0.35 | 0.33 | 0.38 |
| | $\sigma_e^2$ | 0.7 | 0.66 | 0.64 | 0.67 |

**Table 2.**
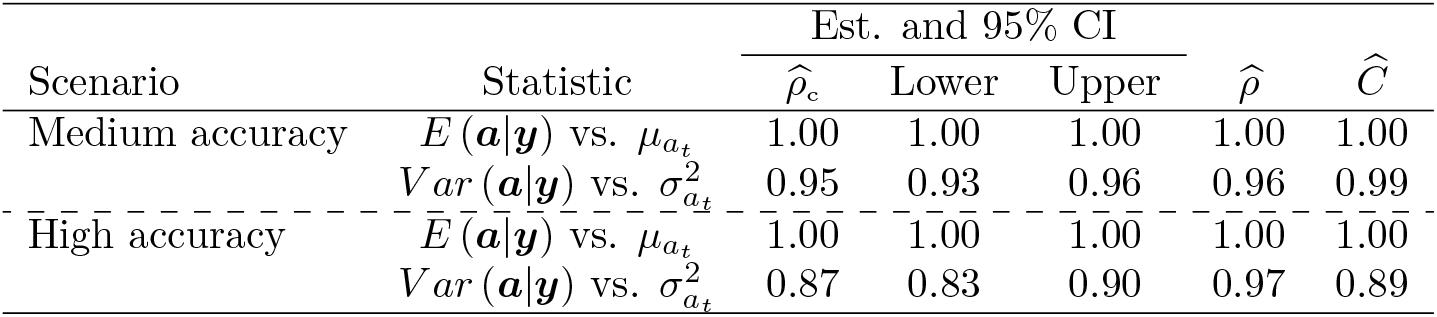
Estimate and 95% confidence interval for the concordance correlation coefficient 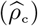 between the estimated and true statistic, and point estimates for the Pearson correlation coefficient 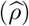 and bias correction factor 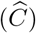 between the variables in each case within scenario

#### 3.2.2 Genetic means and its partitions

Now that the adequacy of the model (7) has been assessed and its impact on the estimates of genetic means and variances over generations has been evaluated, we show the partitioning results when breeding values are estimated from phenotypes. First, we illustrate partitioning results from a single replicate, then we extend it by showing results from 30 replicates. Figure 6 shows the true and estimated genetic mean over 40 generations for the medium- and high-accuracy scenarios considering the total genetic mean (Sum), the path for selected males (M(S)), non-selected males (M(NS)), and females (F). For the medium-accuracy scenario, although the point estimate for the mean of selected males partition showed underestimation), the true means of partition of each path was inside the 95% credible interval. For the high-accuracy scenario, we observed underestimation for females and selected males partition. Consequently, underestimation for the total genetic mean was even higher because it is the sum of those two contributions, while non-selected males had a zero contribution. Figure 7 confirms this result by showing the difference between true and estimated means of partition over 30 replicates. The Supplementary Figure S8 shows that the observed deviations in both scenarios do not come from inadequately estimated Mendelian sampling terms, hence the source of error must be due to the inadequate estimation of the parent average terms.

**Figure 6.**
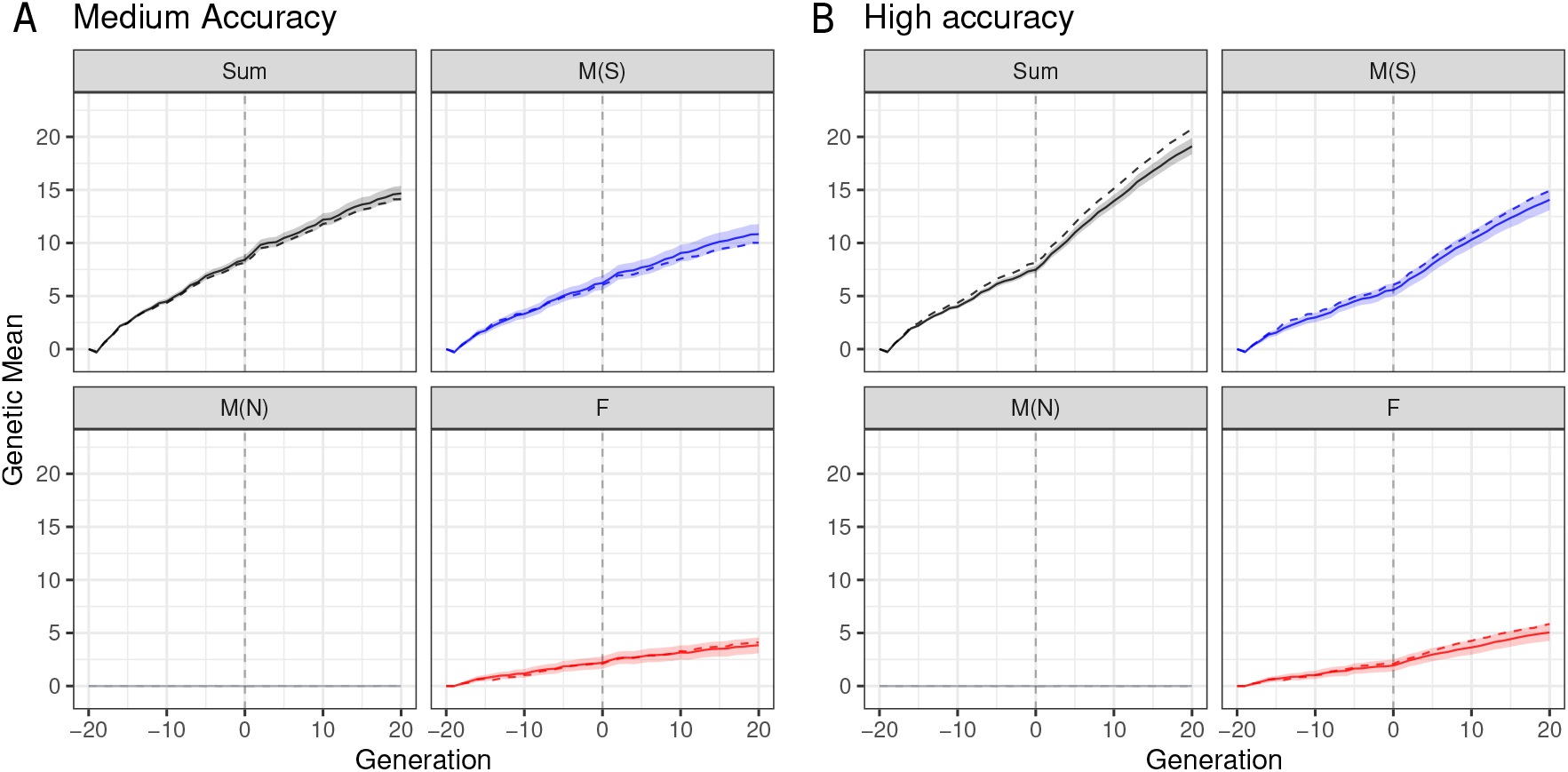
Partitioning of the total genetic mean (Sum) over generations by selected males (M(S)), non-selected males (M(N)), and females (F) paths in the medium-accuracy (A) and high-accuracy (B) selection scenario, considering one replicate (true value is denoted with a dashed line and posterior mean is denoted with a solid line and 95% credible interval is denoted with a ribbon)

**Figure 7.**
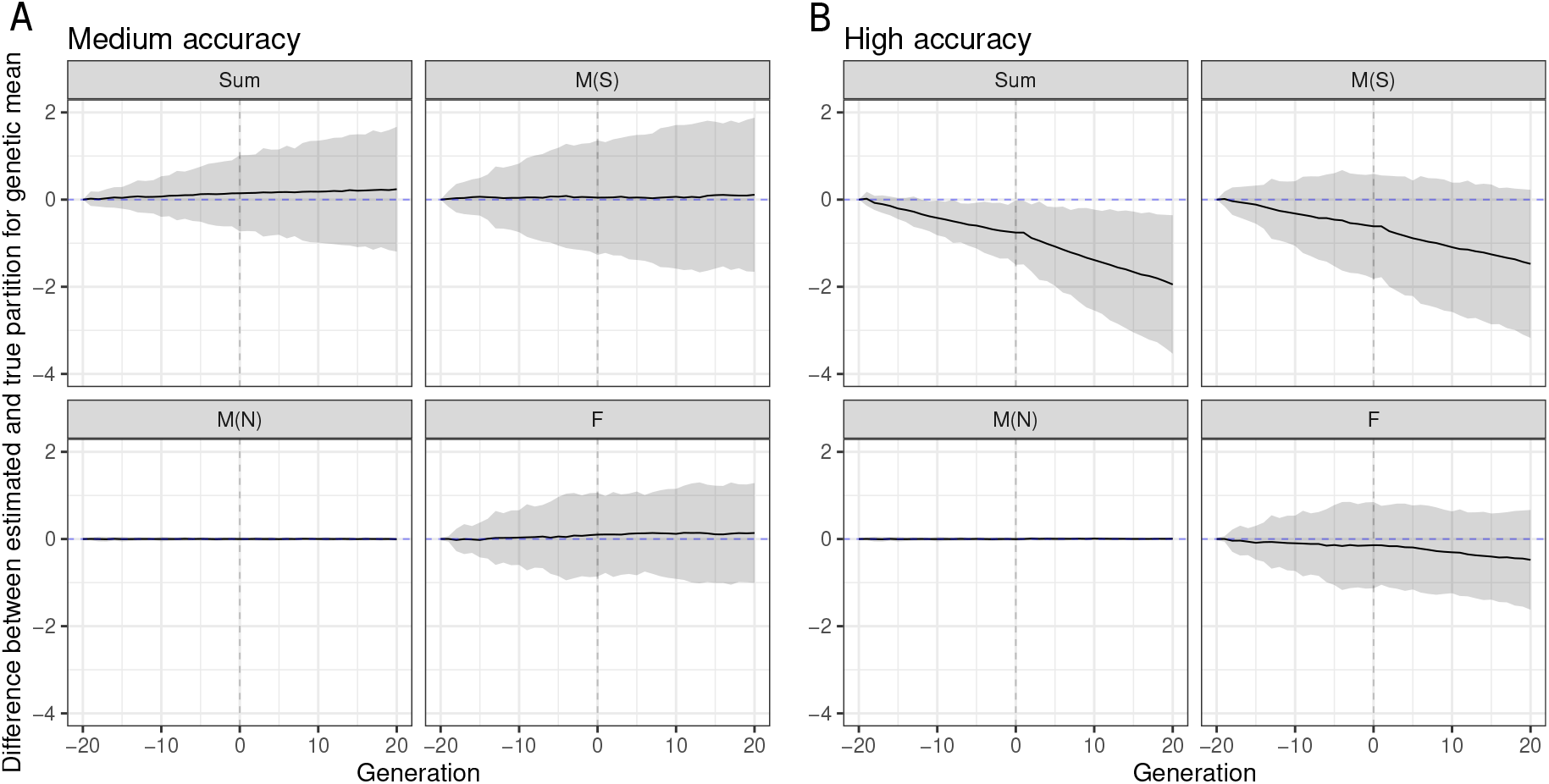
Distribution of the difference between true and estimated genetic means over generations for the total (Sum) partitioned by selected males (M(S)), non-selected males (M(N)), and females (F) paths in the medium-accuracy (A) and high-accuracy (B) selection scenario, considering 30 replicates (zero value is denoted with a dashed line and mean difference over replicates is denoted with a solid line and 95% quantile of differences over replicates is denoted with a ribbon)

#### 3.2.3 Genetic variance and its partitions

The partitioning of genetic variance by paths in the medium- and high-accuracy scenarios in a single replicate are shown in Figure 8. While we correctly estimated the overall trends in the total genetic variance and its partitions, we observed a slight overestimation for the females and non-selected males paths and the total in either the medium- or high-accuracy scenarios. However, from generation 1 to 20 in the high-accuracy scenario, the overestimation increased compared to the medium-accuracy scenario. These observations were confirmed also across 30 replicates for both scenarios (Figure 9). Importantly, distribution over 30 replicates did not include zero in later generations indicating significant departure of the estimates from the true values. Figure 9 also shows an underestimation of genetic variance for the selected males path in early generations (−19 to 2), which lead to the underestimation of the total genetic variance in the high-accuracy scenario.

**Figure 8.**
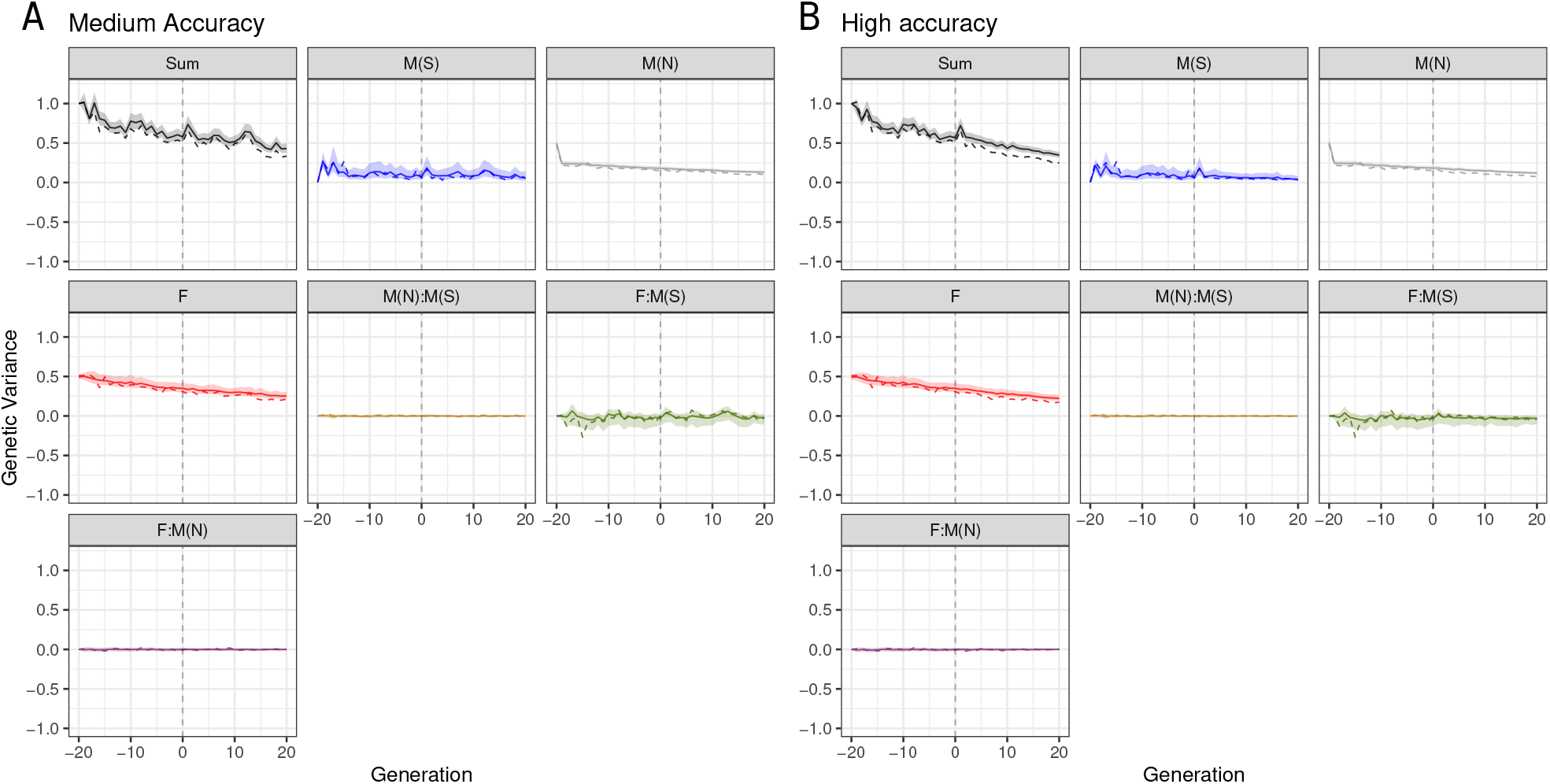
Partitioning of the total genetic variance (Sum) over a generation by selected males (M(S)), non-selected males (M(N)), and females (F) path in the medium-accuracy (A) and high-accuracy (B) selection scenario, considering one replicate (true value is denoted with a dashed line and posterior mean denoted with a solid line and 95% credible interval is denoted with a ribbon)

**Figure 9.**
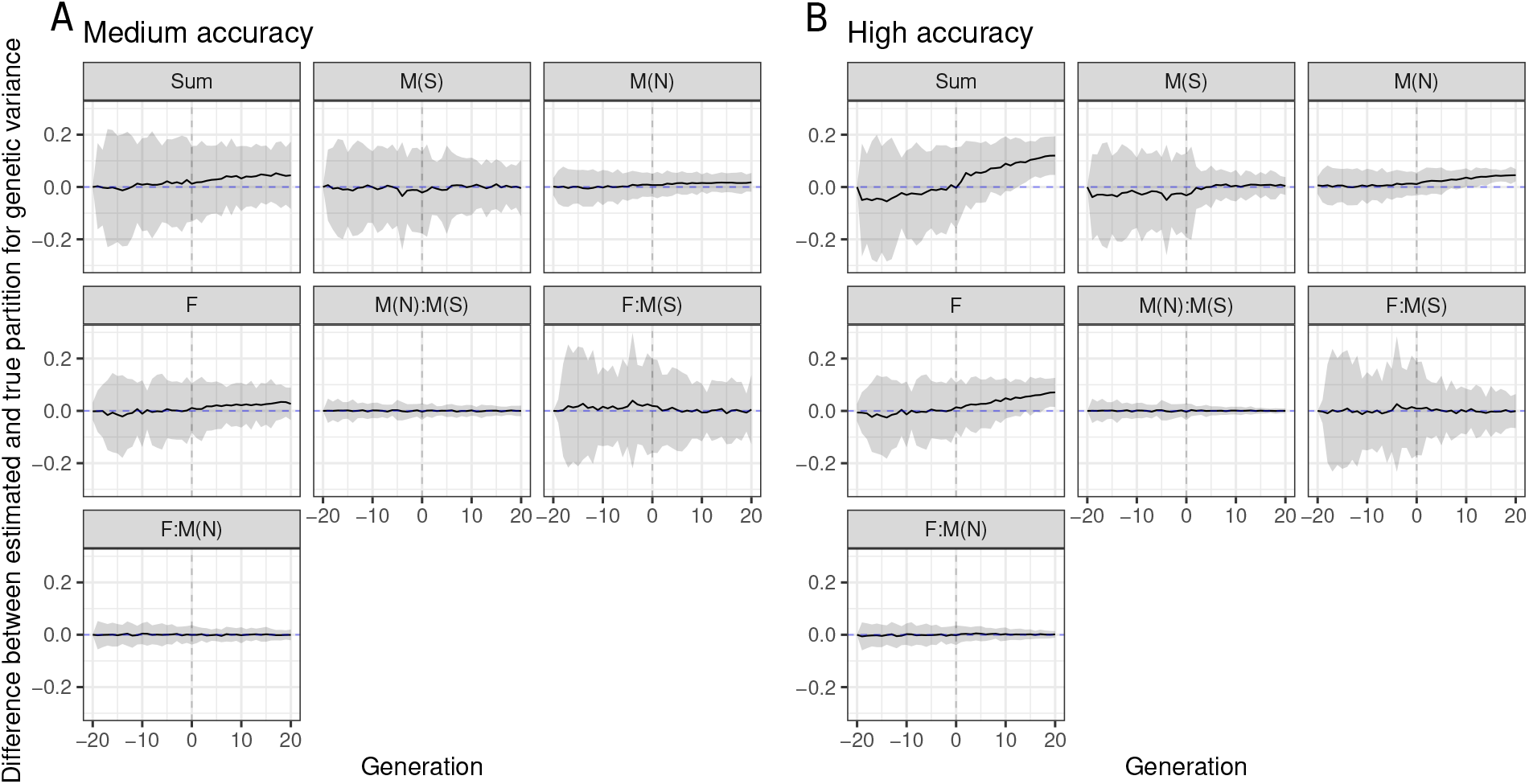
Distribution of the difference between true and estimated genetic variances over generations for the total (Sum) partitioned by selected males (M(S)), non-selected males (M(N)), and females (F) paths in the medium-accuracy (A) and high-accuracy (B) selection scenario, considering 30 replicates (zero value is denoted with a dashed line and mean difference over replicates is denoted with a solid line and 95% quantile of differences over replicates is denoted with a ribbon)

We have initially observed even larger biases but have addressed these by adequately accounting for inbreeding in setting up the ***A***^−1^. Ignoring inbreeding significantly impacted the estimates of genetic means and variances and their partitions (Supplementary Figures S9, S10, and S11).

## 4 Discussion

We developed a method for partitioning the trends in genetic mean and variance into contributions of different paths as an extension of the work of Garcia-Cortes et al. [1] and Obsteter et al. [17]. The method used to infer the path contributions over generations is illustrated using a single-trait model; however, extension to multiple traits is straightforward and already implemented in `AlphaPart`. The extension presented here allows researchers to quantify the drivers of genetic variance in their breeding programmes in addition to the drivers of the genetic mean. Consequently, it could help to understand the dynamics between genetic mean and variance in global animal breeding [2, 4], test how different breeding schemes may impact their long-term sustainability [23, 24], and optimise investment to bring more variability to early stages of the breeding programme [25, 26]. Therefore, it is a powerful and valuable method to understand how different groups of breeding individuals contribute to change in genetic mean and variance, a hot topic that has been discussed in the last few years [5, 7, 27]. For this reason, we implemented this method in the `AlphaPart R` package. The extension has been available since version 0.9.3, freely available from CRAN.

The simulated cattle breeding programme with the medium- and high-accuracy scenarios illustrated the power of the partitioning method to summarise genetic trends in mean and variance. However, some care is needed when using the proposed method. We have shown that the path variable must be considered carefully because a specific choice can lead to a misinterpretation of the contributions, especially regarding the partition of genetic variance. To this end, we recommend plotting the distribution of partitioned breeding values, where partitions can be done with different variables of interest, like sex and selection status in our study.

By partitioning the genetic mean and variance, we showed that in the high-accuracy scenario, the covariance between females and selected males plays an important role when partitioning the genetic variance. Consequently, in this case *V ar* (***a***) *< V ar* (***a***_*F*_) + *V ar* (***a***_*M*_), where *F* and *M* represent the female and male paths. Furthermore, most of the (additive) genetic variance in the breeding programme pertained to females and non-selected males, which were not the most relevant individuals for disseminating genetic gain. A negative correlation between female and selected males partitions in Figure 4 and Supplementary Figure S3 means that the partitions of genetic variance may not be independent. Since the male partition contributes more and more over generations, the female partition has to contribute less, hence the negative correlation between them. In this sense, we demonstrated that the choice of paths is essential and that partitions are not necessarily independent; therefore, they should not be analysed in isolation from each other.

We observed some bias in our estimates of partitions, but we showed that this bias came from the model specification. Hence, the bias is closely related to estimating variance components in populations under selection and low effective population size [28, 29]. Furthermore, the impact of variance component estimation on estimating temporal genetic variances from the model for the high-accuracy scenario was demonstrated by showing a lower concordance correlation between true (*V ar* (***a***)) and estimated (*V ar* (***a***|***y***)) compared to the medium-accuracy scenario. However, this extreme example is essential to show how the bias in variance components affects the proposed partitioning methodology.

This overestimation of the genetic variance and its partitions in the high-accuracy scenario is likely impacted by the “miss-specified” pedigree-based model for such an intense selection and low effective population size (Ne *<* 20) simulated in our study [12]. Namely, we have observed significant changes in the genetic variance of up to 75% over 40 generations. While the pedigree-based model can account for selection [28, 29], it can not account appropriately for such a significant change in genetic variance [6, 7, 12]. Therefore, our next step is to develop an extension of the partitioning method considering genomic data to overcome the issue of working with the expected probability of identity by descent from pedigrees by using the realised identity by descent or state from genomic data [30, 31]. We have recently already extended the Sorensen et al. [6] method for temporal estimation of genetic variance with a pedigree-based model to work with genomic data. Extending the partitioning method of Garcia-Cortes et al. [1] and current work is a natural next step.

## 5 Conclusions

We developed a method to quantify the sources of genetic variance in breeding programmes by partitioning the genetic variance by analyst-defined paths. The developed method can help breeders and researchers better understand changes in genetic gain and variance in specific breeding programmes. Furthermore, the method can be easily applied to real data by leveraging established software to draw posterior breeding values samples given the observed phenotype data. Working with the posterior sample of breeding values also enables straightforward uncertainty quantification in evaluated genetic variance and its partitions.

By partitioning the genetic variance in a simulated cattle breeding programme, we showed the contribution of different groups of individuals (paths) to the variance and that the covariance between paths can also have a substantial contribution. Hence, to comprehend and manage changes in the genetic variance in a breeding programme, we should not consider the contribution of different paths in isolation but should perform a holistic analysis of partitions of the genetic variance.

We observed some overestimating genetic variance and its partitions, but this was caused by the extreme selection in our simulation study and the pedigree-based model, which could not cope with the genetic change both in mean and genetic variance. Our future research will extend the proposed method using genomic data to overcome the model miss-specification under such extreme selection settings.

The developed method for partitioning genetic variance is a powerful and valuable method for understanding how different paths contribute to changes in genetic variance over time, how they interact within a breeding programme, and how they can be optimised.

## Abbreviations

CCC: concordance correlation coefficient
CI: confidence interval
HPD: highest posterior density
MCMC: Markov chain Monte Carlo
REML: residual maximum likelihood
TBV: true breeding values

## Availability of data and materials

Project name: `AlphaPart`.

Project home page: https://cran.r-project.org/package=AlphaPart.

Operating system(s): Windows, MacOS, Linux.

Programming language: R & C++.

Licence: GPL-3.

Data and Code: https://github.com/HighlanderLab/toliveira_alphapart_variance

## Ethics approval and consent to participate

Not applicable.

## Consent for publication

All authors read and approved the publication of the final manuscript.

## Authors’ contributions

GG initiated and supervised the project. TO extended AlphaPart, simulated and analysed the data, and drafted the manuscript. JO, IP, NH, and GG contributed to the discussion of results and revised the manuscript.

## Supporting Information

**Figure S1.**
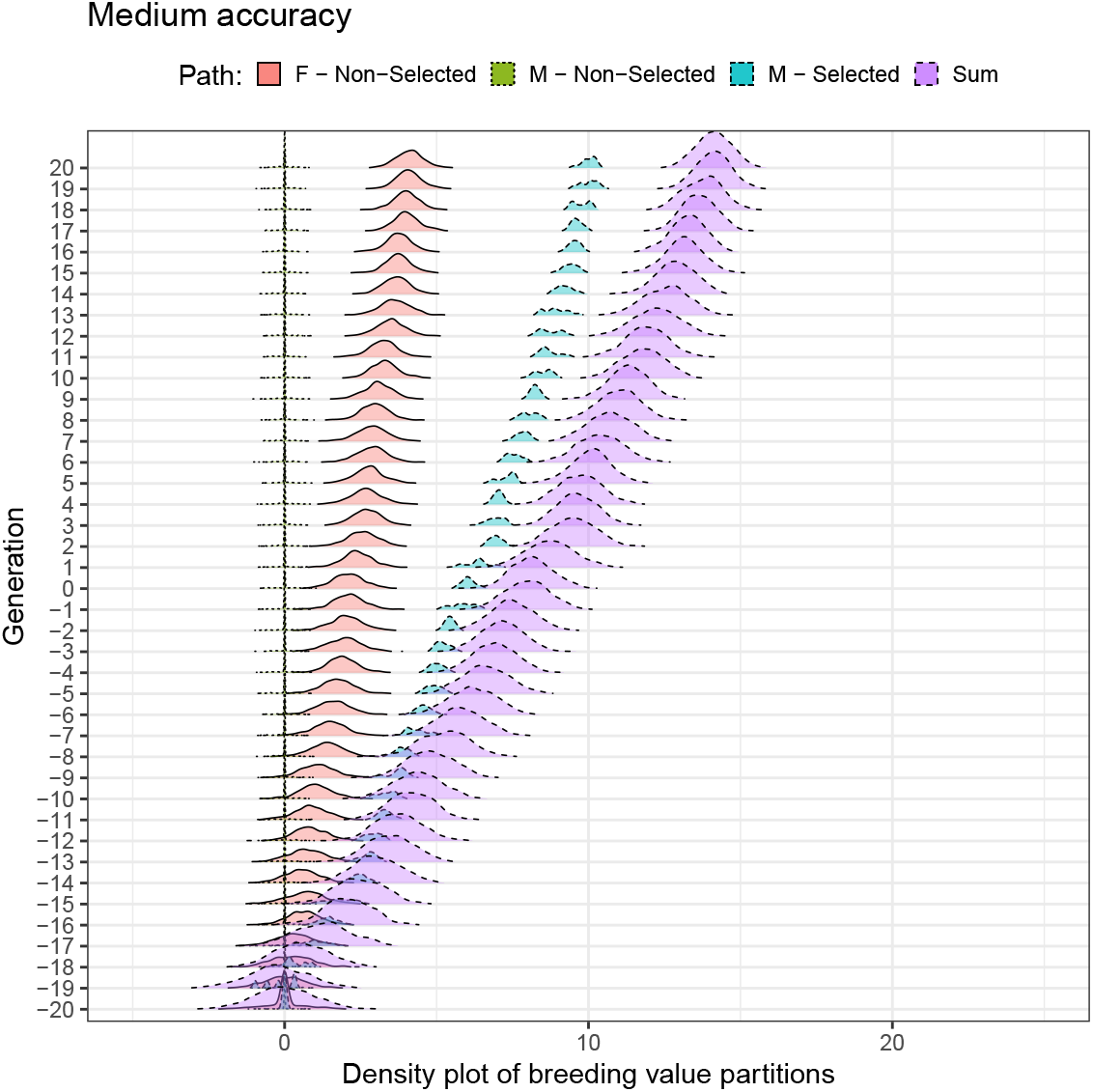
Distribution of breeding value partitions by sex and selection status (selected males (M(S)), non-selected males (M(N)), and females (F)) over generations for medium-accuracy scenario

**Figure S2.**
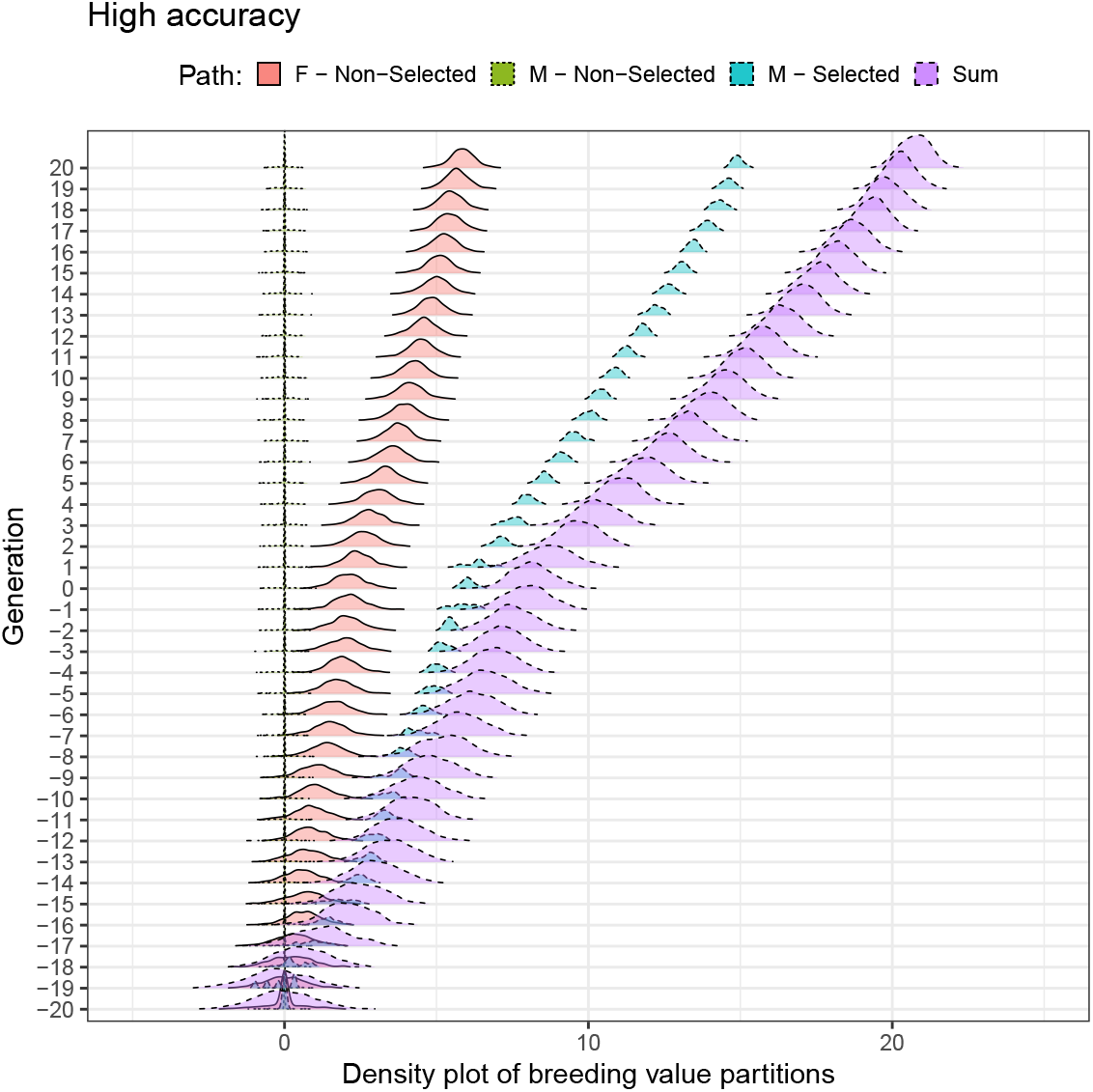
Distribution of breeding value partitions by sex and selection status (selected males (M(S)), non-selected males (M(N)), and females (F)) over generations for high-accuracy scenario

**Figure S3.**
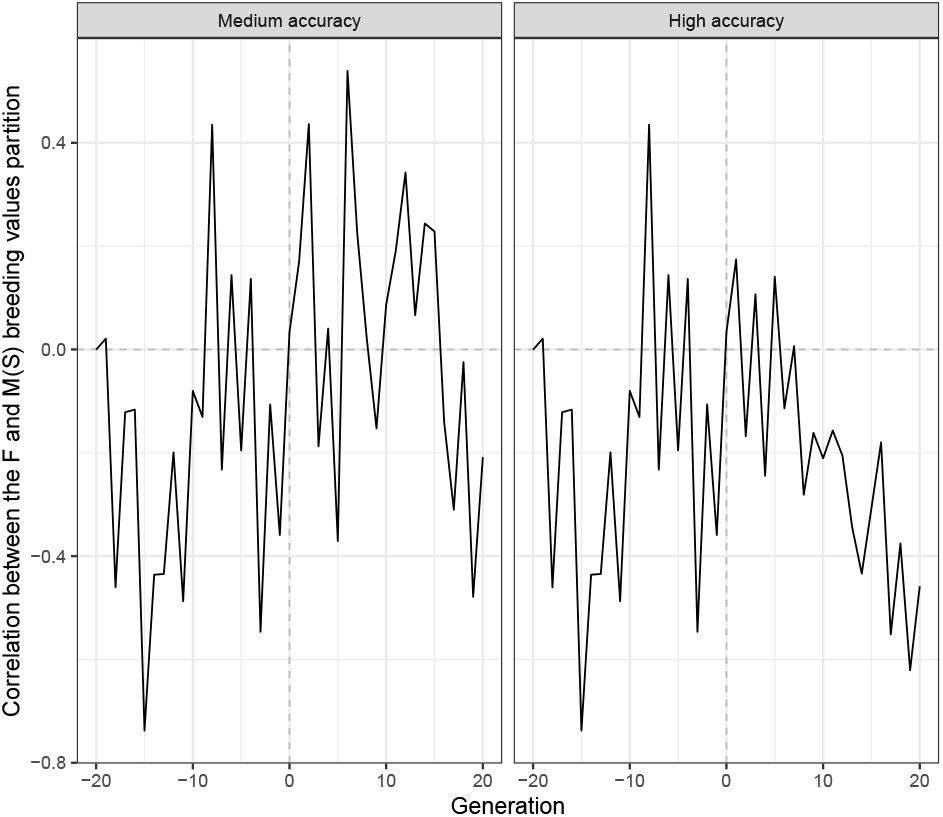
Correlation between Females (F) and selected males (M(S)) partitions using true breeding values for the medium- and high-accuracy scenarios and one simulation replicate

**Figure S4.**
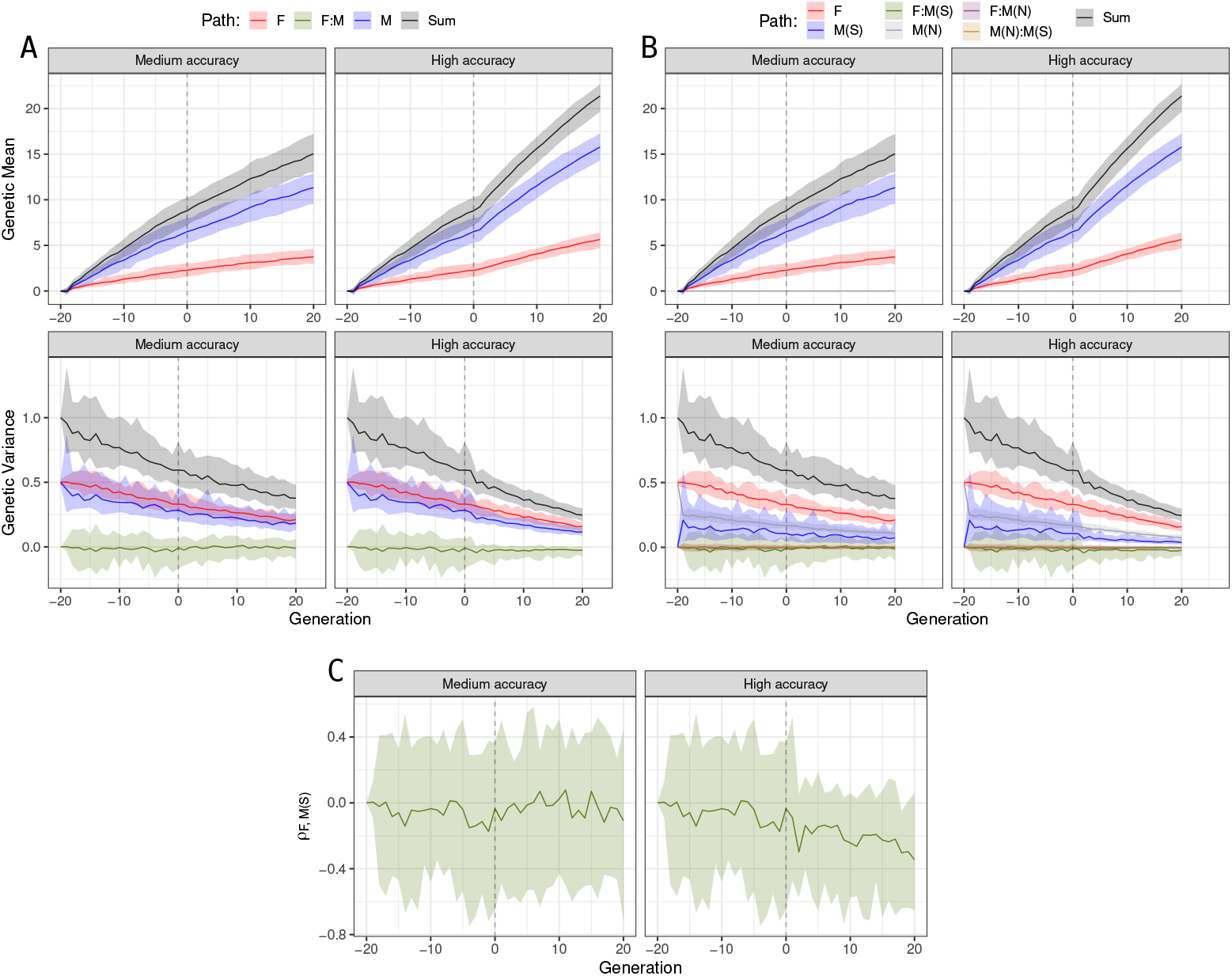
Partitions of genetic mean and variance by (A) sex (males and females), (B) by sex and selection status (selected males (M(S)), non-selected males (M(N)), and females (F)), and (C) the Pearson correlation between F and M(S) partitions for the medium- and high-accuracy scenarios by sex and selection status using true breeding values for 30 simulation replicate

**Figure S5.**
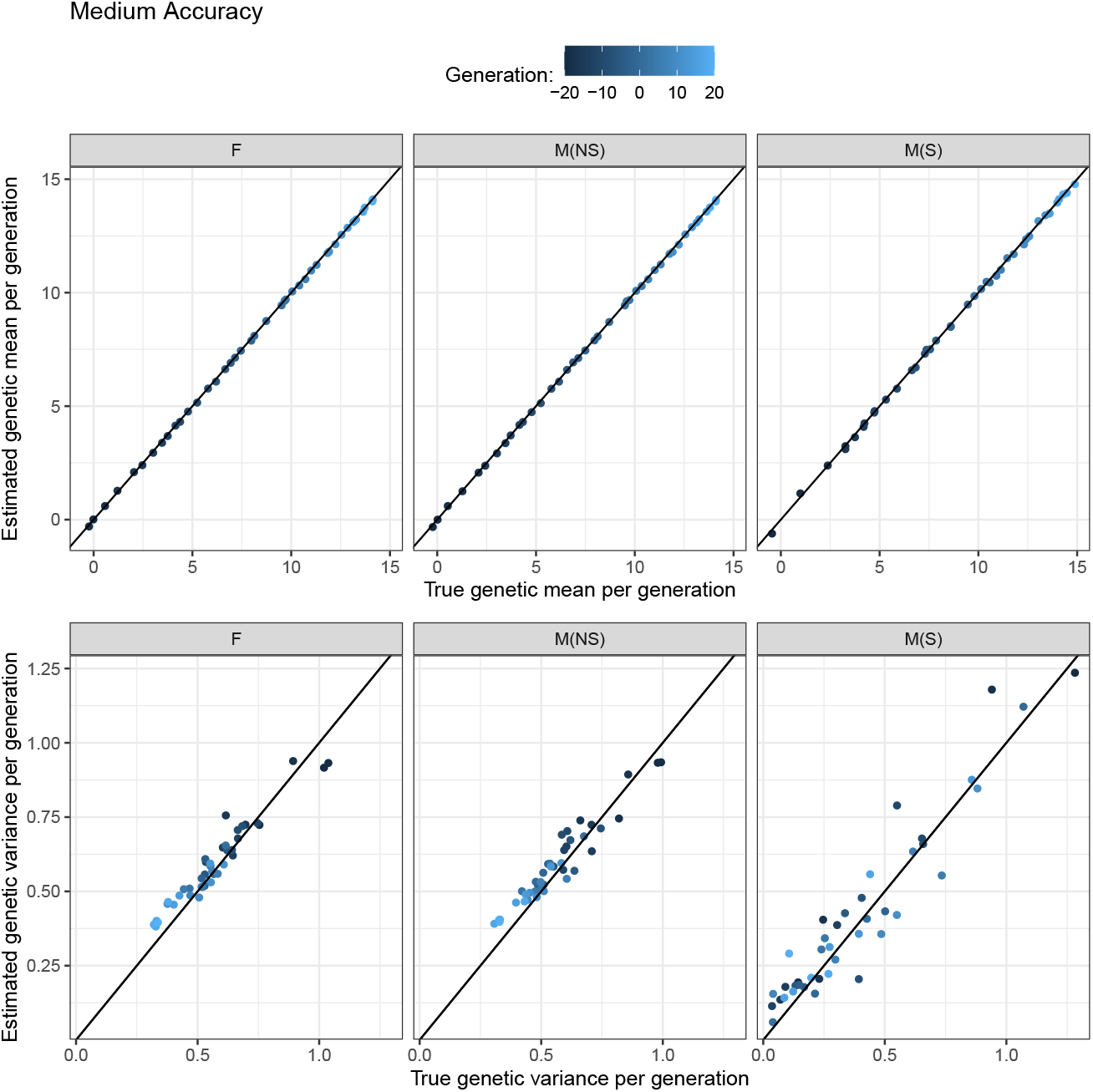
Estimated and true genetic means and variances over 40 generations by selected males (M(S)), non-selected males (M(N)), and females (F) in the medium-accuracy scenario. The solid line represents the equality line *y* = *x* and the dots the Cartesian coordinates of estimated and true values.

**Figure S6.**
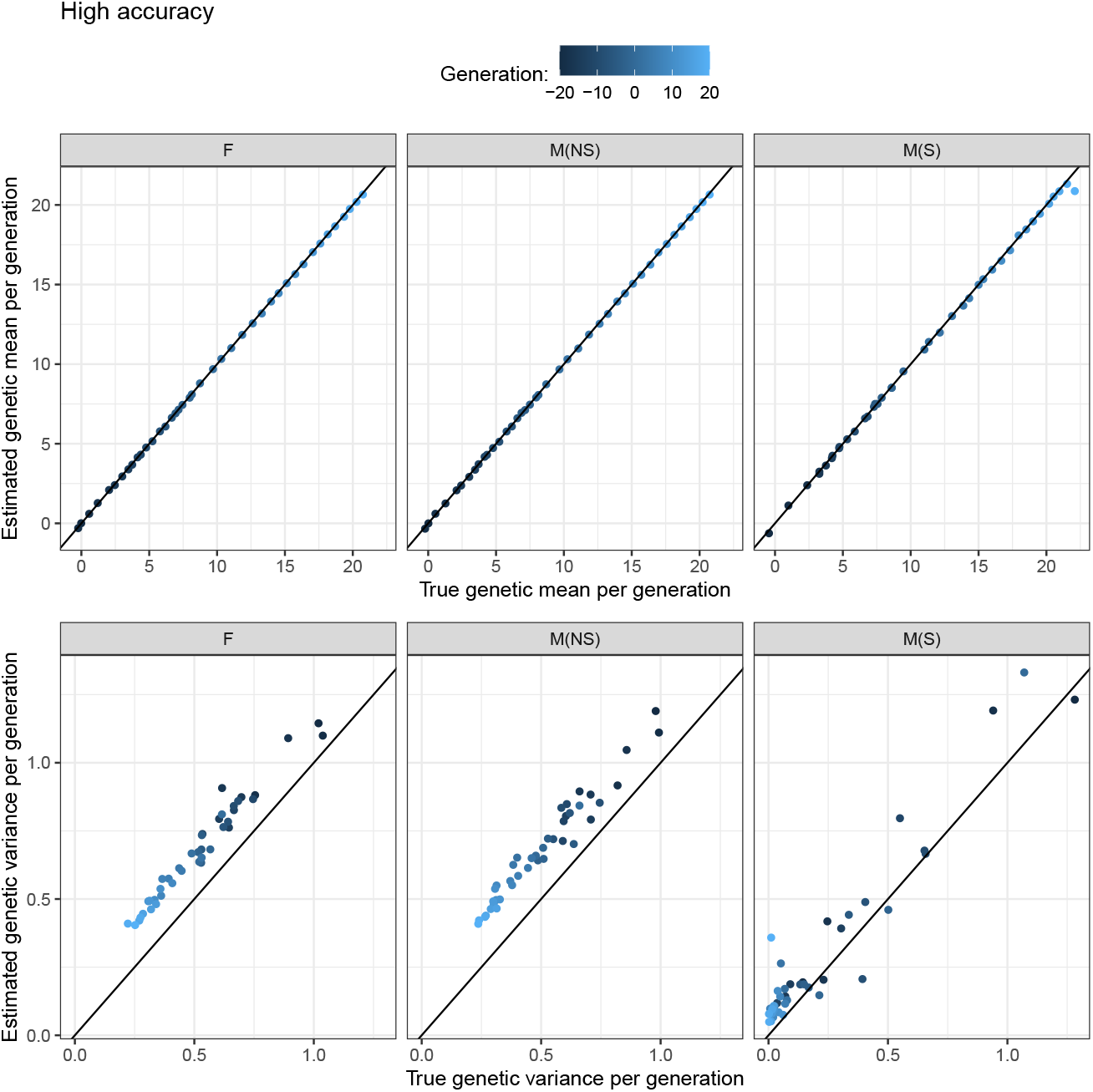
Estimated and true genetic means and variances over 40 generations by selected males (M(S)), non-selected males (M(N)), and females (F) in the high-accuracy scenario. The solid line represents the equality line *y* = *x* and the dots the Cartesian coordinates of estimated and true values.

**Figure S7.**
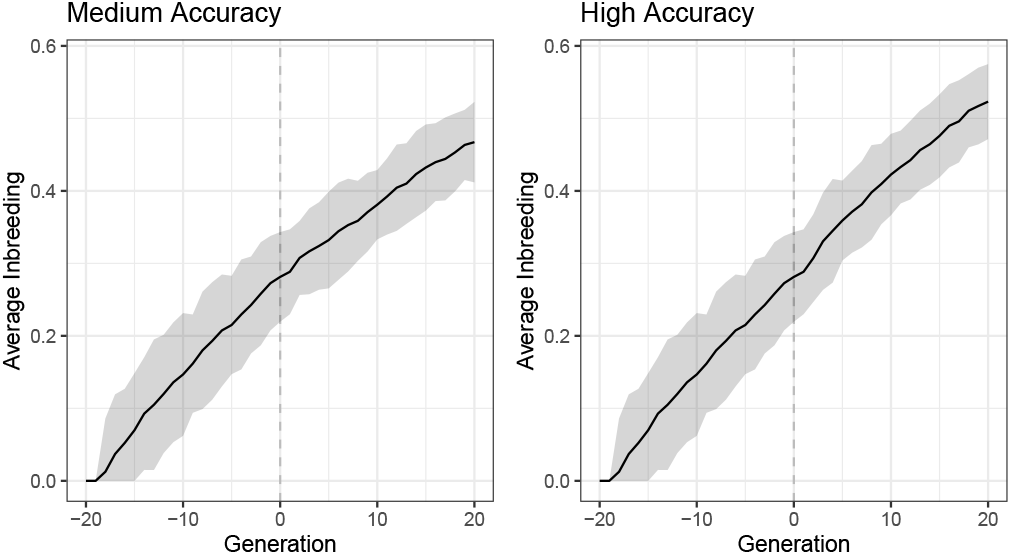
Point (solid line) and interval (ribbon) estimates for inbreeding over generation considering all animals in a specific generation

**Figure S8.**
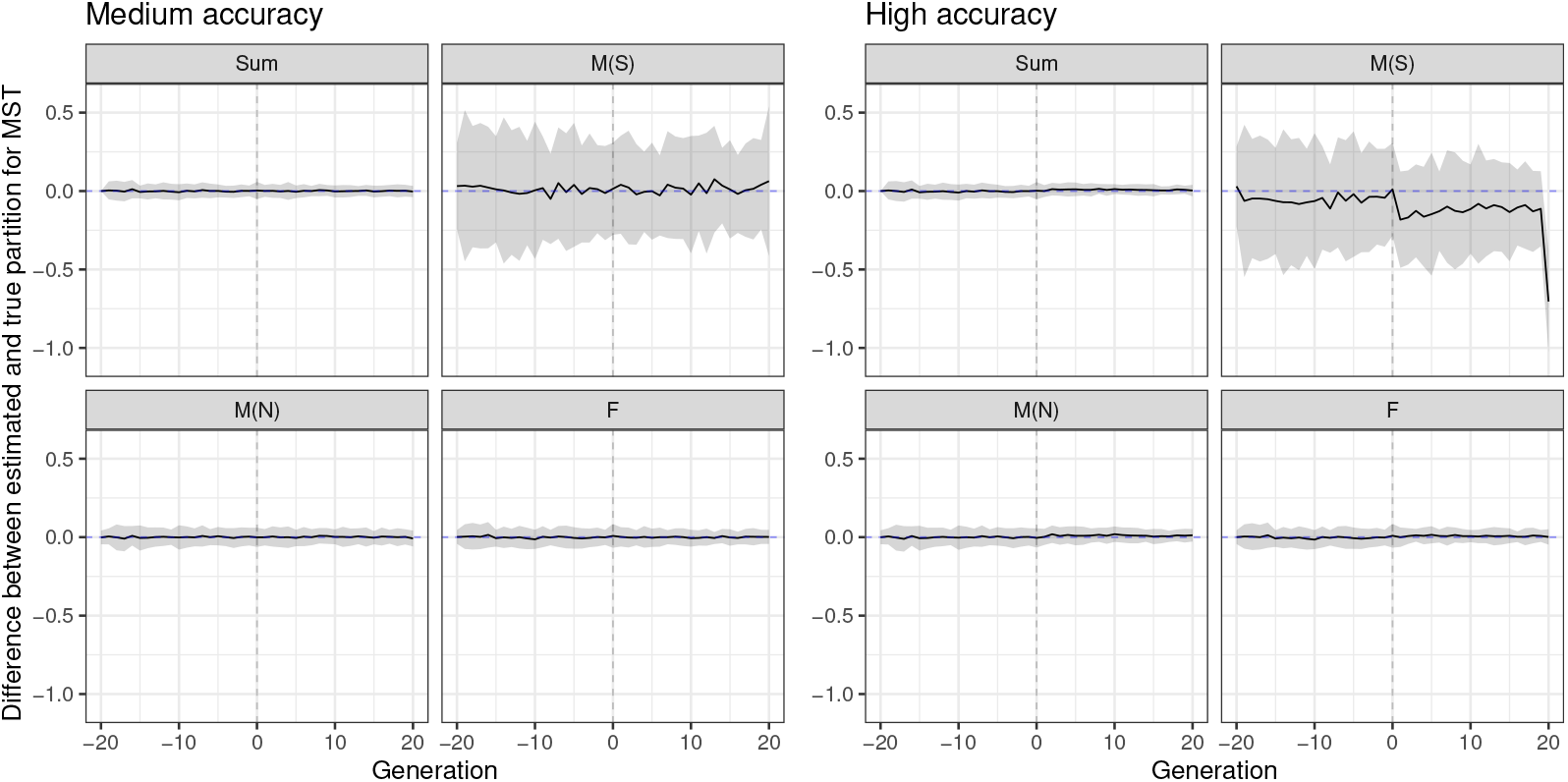
The difference between true and estimated Mendelian sampling terms (MST) is distributed over generations. The total (Sum) is partitioned by selected males (M(S)), non-selected males (M(N)), and females (F) paths in the medium- and high-accuracy selection scenario. We are considering 30 replicates (zero value is denoted with a dashed line and mean difference over replicates is denoted with a solid line, and 95% quantile of differences over replicates is denoted with a ribbon)

**Figure S9.**
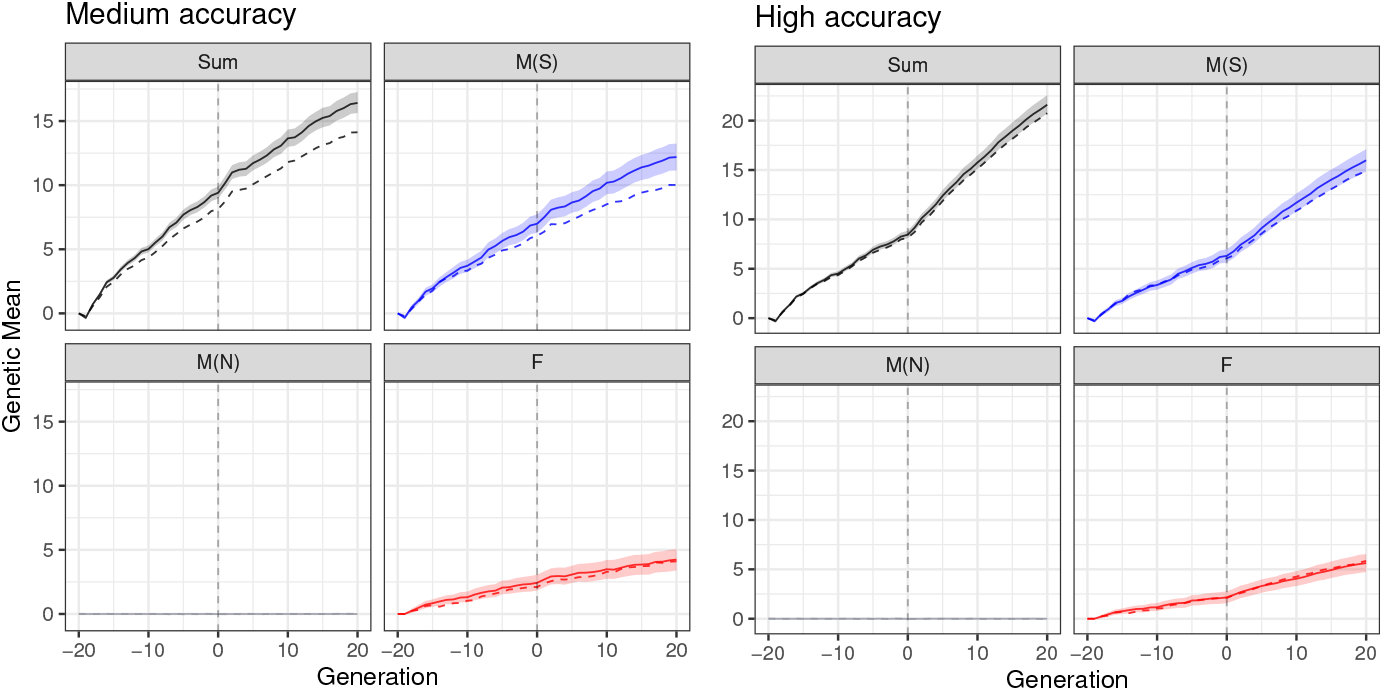
Partitioning of the total genetic mean (Sum) over generations by selected males (M(S)), non-selected males (M(N)), and females (F) paths in the medium-accuracy (A) and high-accuracy (B) scenario. We considered one replicate and without accounting for inbreeding in the model (true value is denoted with a dashed line and posterior mean is denoted with a solid line and 95% credible interval is denoted with a ribbon)

**Figure S10.**
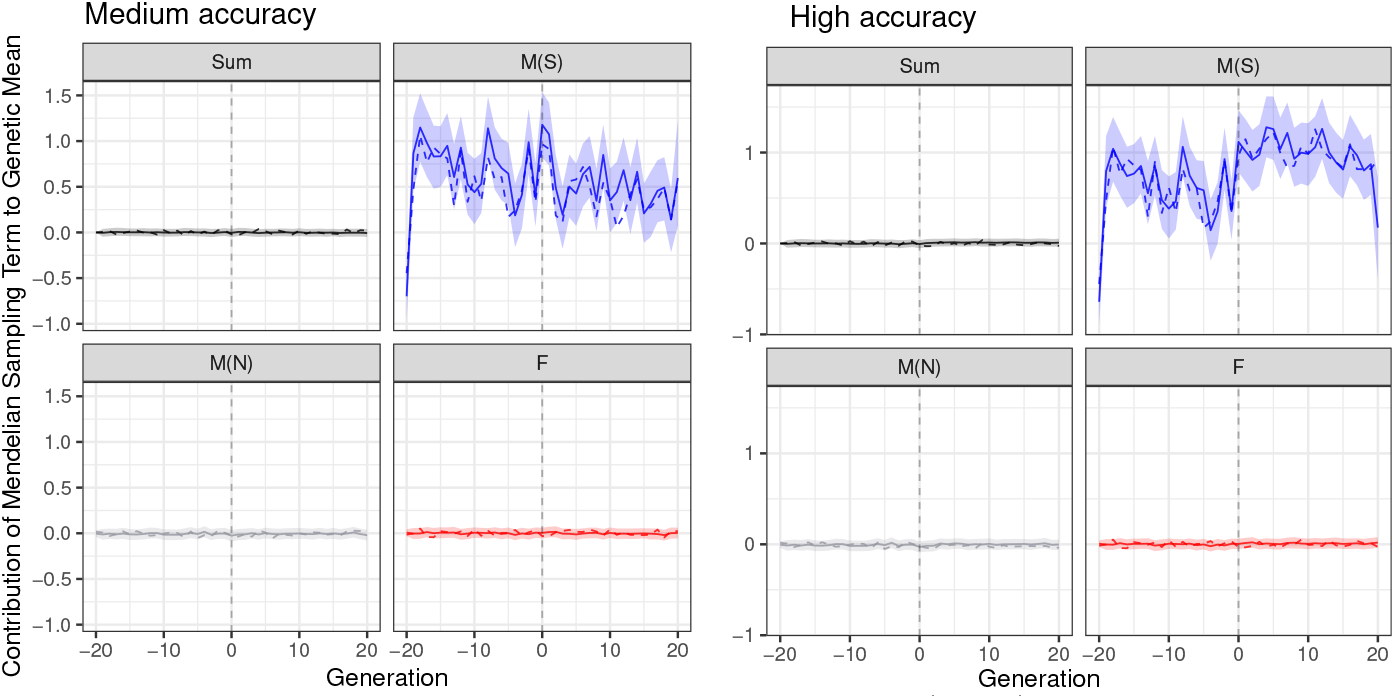
Partitioning of the total Mendelian Sampling term (Sum) over generations by selected males (M(S)), non-selected males (M(N)), and females (F) paths in the medium-accuracy (A) and high-accuracy (B) scenario. We considered one replicate and without accounting for inbreeding in the model (true value is denoted with a dashed line and posterior mean is denoted with a solid line and 95% credible interval is denoted with a ribbon)

**Figure S11.**
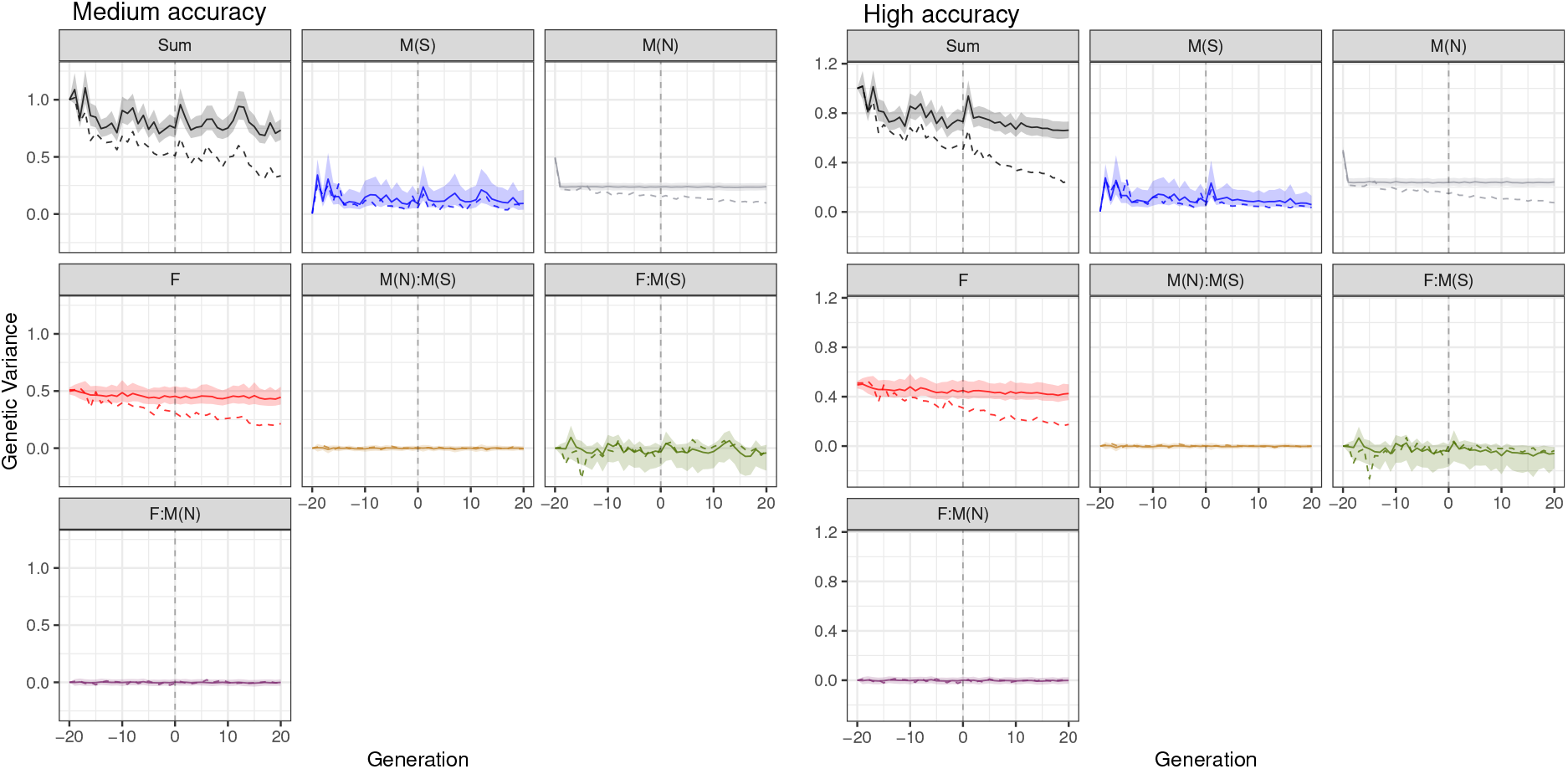
Partitioning of the total genetic variance (Sum) over a generation by selected males (M(S)), non-selected males (M(N)), and females (F) path in the medium-accuracy (A) and high-accuracy (B) scenario. We considered one replicate and without accounting for inbreeding in the model (true value is denoted with a dashed line and posterior mean is denoted with a solid line and 95% credible interval is denoted with a ribbon)

